# A comparative approach for selecting orthologous candidate genes in genome-wide association studies across multiple species

**DOI:** 10.1101/2023.10.05.561051

**Authors:** Lauren Whitt, Elizabeth H. Mahood, Greg Ziegler, Collin Luebbert, Jason D. Gillman, Gareth J. Norton, Adam H. Price, David E. Salt, Brian P. Dilkes, Ivan Baxter

**Affiliations:** Donald Danforth Plant Science Center, St. Louis, Missouri, United States of America; USDA-ARS Plant Genetics Research Unit, Columbia, Missouri, United State of America; School of Biological Sciences, University of Aberdeen, Aberdeen, United Kingdom; Future Food Beacon of Excellence and School of Biosciences, University of Nottingham, Nottingham, United Kingdom; Department of Biochemistry, Purdue University, West Lafayette, Indiana, United States of America

## Abstract

Advances in quantitative genetics have enabled researchers to identify genomic regions associated with changes in phenotype. However, genomic regions can contain hundreds to thousands of genes, and progressing from genomic regions to causative genes is still challenging. In genome-wide association studies (GWAS) measuring elemental accumulation (ionomic) traits, only 5% of loci overlap genes known to control the ionome - indicating that many causal genes are still unknown. To identify candidates for the remaining 95% loci, we developed a method that uses GWAS studies across multiple species to identify conserved causative genes. By Filtering the Results of Multi-species, Analogous, GWAS Experiments (FiREMAGE) we were able to take the GWAS of 19 ionomic traits in Arabidopsis, soybean, rice, maize, and sorghum, and identify alleles affecting trait variation at conserved genes. Permutation testing demonstrated that GWAS loci affecting the same trait contained homologs more often than expected by chance. Most of the top 10% most significant conserved candidate sets encoded alleles in all five species, highlighting the conservation of ionomic genetic regulators in flowering plants. The candidates include proteins with known biochemical functions that regulate the ionome, validating the approach. In addition to genes with known functions, this approach also identified many conserved genes underlying GWAS loci affecting the same trait in multiple species that have no previously identified function in regulating the ionome, providing a path to discover the as yet unknown mechanisms of element accumulation in plants. This method enables the identification of conserved genes of previously unknown function via GWAS.

**Author summary:** Quantitative genetics identifies a genomic region of interest but not the causal gene. We developed an approach to narrow these gene lists using genetic loci affecting elemental (i.e., calcium, iron, zinc) accumulation. Comparative genomics and GWAS demonstrates that alleles at evolutionarily conserved genes alter the same phenotype in multiple species. This produced a list of conserved candidate genes including previously known elemental regulators and genes whose elemental accumulation mechanism has yet to be determined. Combining datasets across species boosted the signal at these loci. This approach accelerates the discovery of new functional roles of genes.

## Introduction

High-throughput phenotyping of elemental accumulation (ionomics) paired with genome-wide association studies (GWAS) has identified many genetic loci associated with variation in the accumulation of elements in plants [1–5]. However, identifying the causal gene among the group of genes linked to the phenotypically-relevant variants, often called a GWAS locus, is challenging for any trait or organism. Typically, many genes are linked to a GWAS locus. Thus each GWAS results in long lists of genes that are equally-weighted candidates for affecting variation in the phenotype. This is especially true for species with large linkage disequilibrium ranges. This has limited the molecular insights provided by studies of natural variation. Complex loci, multiple alleles, rare alleles, incomplete variant information due to a diversity of coding capacity across the species (aka pangenome variation), and sparse marker data further complicate the assignment of causality and can result in the most statistically significant variant being encoded at a position displaced from the causal gene [6]. Additional, and complementary, information about allele consequences and gene function are necessary to identify the molecular cause of detected quantitative genetic variation. Often this relies on labor- and resource-intensive experiments using induced alleles and single-gene testing to provide information of gene functions at candidate loci. Bridging this trait-to-gene gap is necessary to deliver molecular mechanistic insights from studies of natural variation and understand the molecular basis of diversity and adaptation.

Current gene selection methods include examining the functional annotations of genes in a region. Gene ontologies (GO) and annotations derive from our current understanding of known (or predicted) gene functions but are prone to human biases [7,8] and oversights from automatic computational algorithms [9]. GO terms are largely uninformative for novel or under-described traits, including some elemental accumulation phenotypes. GO terms require ongoing curation and submission of newly discovered gene functions from researchers to maintain relevance and utility. Known genes linked to an association are more likely to be selected for further studies [7,10] which produces a form of positive publication bias that reduces the discovery of new gene functions. Additionally, these methods are most effective in species with the most annotations, like *Arabidopsis thaliana* (Arabidopsis), with well-annotated phenotypes. Exploration of candidate genes in a region is less common, and typically results when GWAS identifies a locus with large trait effects. Several approaches have been used to identify causal genes from genes of unknown function. Guided methods that can work without gene annotations include machine learning techniques like PILGRM [15], GenAMap [16], network analysis, gene expression experiments, or a combination of techniques [17]. These methods rely on extensive computation and resources like gene expression data. As a result, they are most valuable in model systems with large collections of preexisting data. Model systems like Arabidopsis also benefit from short generation times and small growth requirements enabling complex and multigenerational designs [18].

Distinguishing likely causative genes among many linked candidates without relying on existing annotations, ontologies, or preexisting data is significantly more complex and time consuming. For example, linkage mapping in experimental line cross populations has been used to differentiate causal candidates following trait-associated GWAS [11,12]. Fine mapping to find a causal gene is expensive and laborious and is not feasible for alleles with moderate or small effect sizes [13,14]. Candidate gene testing by phenotyping knockout and overexpression mutants is an alternative approach and the identification of additional alleles with impacts on the same phenotype is seen as evidence of the correct specification of causal loci. This is facilitated by large collections of mutants, only available for the best resourced model systems such as Arabidopsis and *Zea mays* (maize) with substantial mutant resources. Gene editing has made these methods less dependent on pre-existing populations and expanded the possibilities of candidate gene testing outside of traditional model systems. The facility of these experiments is still dependent on generation time and tissue culture, transformation, and plant regeneration. This makes the identification and prioritization of short lists of candidate genes essential for feasibly testing candidates.

Developments in whole genome and next-generation sequencing have improved *de novo* genome assembly of complex genomes and the identification of genetic markers (i.e., SNPs, indels) within populations [19]. This makes it conceivable that comparison of alleles discovered in multiple species might form the basis of validation experiments. When independent alleles in homologs have similar phenotypic impacts, this is functionally equivalent to annotation of function by inference from sequence similarity. Many agriculturally important crop species have excellent genome assemblies and annotations. Arabidopsis still has the highest quality plant genome, annotations, and resources that inform annotations in other organisms. For some agricultural species (e.g., soybean), gene annotations are inferred directly by homology searches against Arabidopsis. Seeking gene functions using resources beyond arabidopsis can dramatically enhance our understanding of plant biology, including the biology of crops.

The ionome describes the composition of elements in plant tissue, all of which is acquired from its environment [20]. Obtaining, excluding, and homeostatically maintaining the concentration of elements in cells are functions shared by organisms across all domains of life. Elements obtained from the environment are critical cofactors and substrates for multiple biochemical reactions, and solving the challenges of balancing the ionome in cells was a necessary step in the evolution of cellular life. As a result, many of the mechanisms that achieve elemental homeostasis are ancient, and experimental approaches for mechanistic validation, such as cross-kingdom and cross-domain molecular complementation (e.g., complementation of bacterial or yeast mutants with plant genes), are highly effective [21,22]. Moreover, as elements are the same regardless of which species they are measured in, we can directly compare elemental accumulation traits across species. As a result, the field has repeatedly demonstrated that natural alleles at homologous genes affect the levels of the same elements in multiple plants [22–31]. This independent evolution of alleles with similar phenotypic consequences in multiple species is the hallmark of conserved critical regulators. The direct comparability of ionomics traits, repeated observation of natural variation in homologs affecting phenotypic variation, and the deep conservation of mechanisms affecting its variation suggests that we could use homology to identify the causative genes for ionomic trait variation within GWAS loci across multiple species.

We created a computational method to take the genes linked to GWAS data sets from multiple species and detect orthologous genes present among the linked candidates. To assess the statistical significance of the homology overlaps, and determine if orthology occurred more often than expected by chance, we built a permutation-based assessment of the effectiveness of this tool that calculates the expected background distribution of orthologs contained in a set of genomic regions drawn from multiple species. Comparison of permuted sets to observed orthology overlap demonstrated that GWAS detects alleles at orthologous genes more often than expected by chance. This is the expectation if variation in elemental accumulation is accomplished by conserved genes across flowering plants. We validated these results using lists of genes of known ionomic function [26] and demonstrated that our combination of GWAS and orthology detected alleles at genes known to affect the ionome at a higher rate than expected by chance. However, genes of known function were overwhelmingly in the minority and it identified as likely candidates many genes with no previously known functions. This approach, which we call Filtering Results of Multi-species, Analogous, GWAS Experiments (FiREMAGE), identifies candidate genes without regard for prior evidence of function in element accumulation. This frees the process of candidate selection from the constraint of prior demonstration of function and permits gene function discovery through the study of natural variation directly. This approach can be applied at any dataset and encourages exploration of novel gene functions affecting trait natural variation.

## Methods

### Summary of Approach

We created a pipeline to compare GWAS results across species and determine if orthologous candidate genes underlie loci of phenotypic impact. The first step is processing of GWAS results to generate genomic intervals affecting the same trait in each member of a set of species. The genes within these intervals are tabulated and the orthologs of those genes in each member of the species set are determined. Permuted genomic regions, matched to each species’ input, are generated and processed in the same manner as the experimental data. Comparison of the background distribution of ortholog overlap, from the permutations, and the experimental observations are made for each set of species and for every trait.

Briefly:

1. Following a GWAS in species 1, regions containing linked SNPs affecting trait variation are identified and retained as windows. The trait’s top associated SNPs in each retained genomic window are identified and used to uniquely name the GWAS locus and summarize the significance and effect direction of the quantitative trait locus.
2. This is repeated for every species. At this point a set of genomic regions, linked to GWAS hits is established for the trait across all species.
3. Candidate genes are identified as encoded within these experimentally determined windows. At this point, overlaps between the candidates and known ionomic genes can take place (this is separate from FIREMAGE and is a standard approach with GWAS made highly effective by the existence of a curated list of known ionomic genes).
4. Orthology relationships for all candidates from step 3 are determined across the five species using Orthofinder. At this point, the number of GWAS loci containing orthologous candidate genes were determined. These orthologous genes are returned as FIREMAGE candidates. This is the first step that is unique to our FiREMAGE approach.
5. FiREMAGE assesses the statistical significance of the orthology overlaps using permutation testing. 1000 sets of genomic regions with size-matched windows, from these five genomes, are selected at random. The rate at which these randomly permuted regions return overlapping orthologs is then used as the expected distribution of orthology overlaps for each trait given the observed number and size of genomic regions across all five species. This is a permutation test of the expected number of overlaps for windowed regions of the sizes returned from the GWAS for each species.
6. The False Discovery Rate of GWAS candidate gene ortholog overlaps is determined by comparing the true number of orthologous genes among the genomic regions affecting a trait across these species (Step 4) to the number determined for a permuted set of genomic regions across these species (Step 5).
7. This pipeline is repeated for every trait.

### Data availability

FiReMAGE scripts, GWAS SNPs used in this paper, ortholog tables, and gene coordinates can be found at https://github.com/danforthcenter/FiReMAGE. Gene coordinates can alternatively be downloaded from Phytozome (https://phytozome-next.jgi.doe.gov/phytomine/begin.do).

R scripts created for this newest version of the Known Ionome Gene (KIG) List used to validate FiReMAGE results can be found at https://github.com/danforthcenter/KIG_v2.

### GWAS datasets

#### Single nucleotide polymorphism (SNP) filtering

We used published elemental profiling GWAS datasets from *Arabidopsis thaliana* (Arabidopsis) [32], *Oryza sativa* (rice) [33–36], *Sorghum bicolor* (sorghum) [1], and *Zea mays* (maize) [3]. The *Glycine max* (soybean) data is described below. The p-value thresholds used are intentionally permissive to reduce false negatives in our GWAS/ortholog overlaps. False positive control is provided by requiring orthologous genes to exceed multiple thresholds and estimated by permutation, see Random Permutation, below. The SNPs from each GWAS dataset were selected by p-value thresholds that were separately determined for each species: *Arabidopsis thaliana* (Arabidopsis; p < 0.001), *Glycine max* (soybean; p < 0.05), *Oryza sativa* (rice; p < 1e-5), *Sorghum bicolor* (sorghum; p < 1e-4), and *Zea mays* (maize; p < 0.01).

The fundamental units that FIREMAGE uses are genomic ranges from each species corresponding to each locus identified by GWAS. To provide SNP-to-locus identification, and delimit the genomic range corresponding to each locus, SNPs from each dataset were collapsed into loci using linkage ranges from each species (Table 1). The SNP with the lowest p-value (e.g. the most significant or “top SNP”) for each range was used to name and annotate each locus. The p-value of the top SNP was retained as the collapsed loci’s p-value. This SNP-to-locus mapping procedure results in no overlapping ranges and avoids any problems associated with genes occupying more than one overlapping locus. All references to “loci” provided to FIREMAGE refer to these genomic intervals and the use of the word “locus” and not “gene” is deliberate as these locations can comprise multiple genes. In addition to GWAS SNP-to-locus mapping, the FIREMAGE pipeline can accept any file that describes loci as genomic intervals. As a result, it is already constructed to accept the outputs of procedures such as joint linkage analyses in nested-association mapping populations such as the maize NAM dataset.

**Table 1.** Overview of GWAS datasets.

| Table 1. Overview of GWAS datasets. |  |  |  |  |
| --- | --- | --- | --- | --- |
| Species Annotation | general linkage range | input as loci | SNPs/loci | collapsed loci |
| *A. thaliana_TAIR10* | 75000 | FALSE | 7167 | 1404 |
| *G. max_Wm82.a2.v1* | 200000 | FALSE | 37854 | 7308 |
| *O. sativa_v7.0* | 243000 | FALSE | 17109 | 432 |
| *S. bicolor_v3.1.1* | 100000 | FALSE | 2139 | 1768 |
| *Z. mays_RefGen_V4* | 100000 | TRUE | 905 | 905 |

#### Soybean GWAS and 2015-2017 field experiments

*Glycine max* genotypes were obtained from the USDA-ARS GRIN collection (https://npgsweb.ars-grin.gov/gringlobal/). Field experiments were performed over three years (2015, 2016, and 2017) with three field replicates at each field experiment in a random complete block design. The 2015 and 2016 field experiments were performed at the University of Missouri Bradford Research farm located near Columbia, Missouri, (38°53’38.4"N 92°11’47.1" W) which has a Mexico silt loam (fine, smectitic, and mesic Vertic Epiaqualf) and the 2017 field experiment was performed at the Rollins Bottom research farm located near Columbia, MO, ((38°55ʹ37.5ʺ N, 92°20ʹ44.6ʺ W) on a Haymond silt loam soil (course-silty, mixed, superactive, mesic Dystric Fluventic Eutrudepts). Seeds were planted at a rate of 8 seed/ft, over 6 ft with a 2 ft gap between plots in 2015 and 2016, and 3 seed/ft over 8 ft with a 4 ft gap between plots in 2017. Row spacing was 30” between rows in all experiments. Field conditions were typical of soybean production in the Midwest USA, with NPK fertilizer applied at rates appropriate per soil analyses, and pre-emergent and post-emergent herbicides were applied as recommended to control weeds for conventional soybeans. Field plots were planted on 06/30/2015, 05/06/2016, and 06/06/2017. Plots were hand-harvested and threshed using a small plot thresher by 10/29/2015, 09/23/2016, and 11/17/2017. Flower color for each field plot was recorded and compared to known colors for each genotype. Similarly, seeds were inspected after threshing and compared to known hila and seedcoat coloration, with non-matching samples discarded. Approximately 25 seeds were lyophilized for each field plot and ground to a fine powder using a Perten 3310 lab mill using a “type 0” grinding disc (Perten Instruments, Sweden). Powdered samples from each grow-out were analyzed concurrently for concentrations of 20 elements using an inductively coupled plasma mass spectrometry (ICP-MS) pipeline previously described [37]. Batch effects inherent in the ICP-MS process were confounded with grow-out in the initial analysis. Thus, a subset of samples from each batch of the initial ICP-MS analysis were re-analyzed for elemental concentrations, and a batch-specific correction factor was generated and subtracted from the first run’s values. Analytical outliers identified as > 10 median absolute deviations away from the median were removed within grow out.

For analysis within each grow-out, phenotypic values for each genotype were taken as the median value within that grow-out. To better meet normality assumptions both for the GWAS and for aggregation of the grow-outs using a mixed model method, phenotypes were Box-Cox transformed within the out and across grow-out and then standardized. Lambda values for the Box-Cox were chosen using a 95% confidence interval. To aggregate data from different grow-outs, best linear unbiased predictors (BLUPs) for each genotype were calculated from a random intercept mixed model: 𝑃ℎ𝑒𝑛𝑜𝑡𝑦𝑝𝑒 ∼ (1|𝐺𝑒𝑛𝑜𝑡𝑦𝑝𝑒) + (1|𝐺𝑟𝑜𝑤 𝑜𝑢𝑡) + (1|𝐺𝑒𝑛𝑜𝑡𝑦𝑝𝑒: 𝐺𝑟𝑜𝑤 𝑜𝑢𝑡).

Association mapping was performed for each phenotype in the three grow-outs and the combined grow-out BLUPs. Genotype sample sizes for each grow-out are 355, 363, and 364 for 2015, 2016, and 2017, respectively. To account for population structure, kinship matrix and principal components were included in the mixed model as fixed effects. The kinship matrix was calculated using all 365 genotypes present in at least one grow-out following the realizedAB method in the synbreed R package [38] as implemented in the R package ionomicsUtils (v1.0 https://github.com/gziegler/ionomicsUtils). Principal components included in the model were selected for each phenotype if they were correlated significantly with that phenotype at the α = .001 level.

The multi-locus mixed model (MLMM) used performs forward inclusion and backwards elimination until all of the heritable variation is explained or a maximum number of cofactors has been reached, which in this study was 40. Model selection was performed using the extended Bayesian information criterion [39] and the multiple-Bonferroni criterion [40]. A third set of SNPs was also returned based on inclusion in the final forward inclusion step (max cofactor). Effect sizes for SNPs returned were calculated using the mlmm.gwas package (S10 Table) [41]. A resulting SNP was said to overlap with a neighboring SNP if the r-^2^ linkage disequilibrium was > 0.2 as calculated using the LD function in the genetics R package [42] and implemented in the R package ionomicsUtils (v1.0 https://github.com/gziegler/ionomicsUtils). Average linkage disequilibrium values for all pairs of SNPs were calculated across the genome by first separating SNP pairs into 100 kB bins based on distance and then calculating LD within these bins [43].

### OrthoFinder orthologs

OrthoFinder v2.0 [44] was downloaded from https://github.com/davidemms/OrthoFinder and used with default parameters and best practices outlined here: https://davidemms.github.io/orthofinder_tutorials/orthofinder-best-practices.html. Protein reads of primary transcripts were obtained from Phytozome (v13) [45]. To make the Known Ionome Gene (KIG) list, we used files for all species in the list except for wheat due to its low-quality genome annotation. *Liriodendron tulipifera* was included as the most closely related outgroup to the eudicots and monocots in Phytozome v13. The resulting ortholog table included one-to-one, one-to-many, and many-to-many ortholog relationships.

### Loci/gene overlaps

We obtained all the Phytozome (v13) [45] gene coordinates for each species via the BioMart tool (v0.7) located on the Phytozome homepage (https://phytozome-next.jgi.doe.gov). Using the coordinates, we search for genes overlapping trait loci that are also in our 5-species ortholog groups. For each trait, FiReMAGE retains genes from ortholog groups with overlaps in at least 3/5 species and produces a list of prioritized candidate genes. The number of overlapping orthologs is recorded for each locus to calculate false discovery rates.

### Random permutations

GWAS loci permutations were 1000 selections of random locations equal to the actual datasets’ species and trait composition. Each species’ chromosome start and end coordinates were downloaded from the European Nucleotide Archive (ENA) database (GCA_000001735, GCA_000004515, GCA_001433935, GCA_000003195, GCA_000005005). Because random base pair locations are unlikely to be in range for collapsing into “loci”, we assigned range lengths randomly chosen from the actual GWAS loci. If random locations are close together, their randomly assigned ranges may overlap. FiReMAGE runs the random permutations through the same loci/gene overlap pipeline as the actual data, calculates the permutation’s distributions to evaluate the likelihood of a locus overlapping any nearby ortholog, and produces a false discovery rate (FDR) to prioritize candidate genes. FDR was calculated as the mean number of orthologs overlapping random locations divided by the number of actual orthologs overlapping loci.

Permutations were also carried out to assess the significance of the overlap with the KIG. Primary KIG list permutations (S3 and S4 Figs, S6 Table) were formed by selecting random genes to replicate the same species and element compositions of the actual primary KIG list and repeating 1000 times. The extended KIG list permutations were formed by putting the primary genes permutations through the same ortholog-finding process as the actual KIG list, with one deviation. In the actual extended lists, a primary KIG’s paralogs (if any) and orthologs are sourced from its ortholog group. In the permutations, we only take the random gene’s orthologs from its ortholog group. After tabulating all orthologs of each set of randomly selected genes in each permutation iteration, we add a random selection of genes from Arabidopsis, rice, and maize that equal the number of paralogs in their actual extended KIG lists. The number of paralogs added to the permutation sets equalled the number of paralogs that exist in the extended KIG list, and does not expand groups where multiple primary genes are also paralogs to each other. This choice was made to avoid random gene selections in which large, single-species, expansions of a gene family filled up the permutation set as this would not correctly replicate the structure of the KIG list and would not do a good job of estimating the background rate of overlap. Known ionomic genes are largely conserved across species, only about 9% of the primary KIG list in the first version of the KIG list were single-species genes [26].

## Results

### Known Ionomic Genes as GWAS candidates

Elemental accumulation, or ionomic, datasets from association mapping populations of five species (Table 2): Arabidopsis [32], *Glycine max* (soybean; see methods), *Oryza sativa* (rice) [33–36], *Sorghum bicolor* (sorghum) [1], and maize [3] were reanalyzed to identify alleles at conserved orthologous genes. These GWAS data sets were selected because they are publicly available and contain the same 19 elemental accumulation traits, measured with the same type of instrument (ICP-MS), and all measurements were obtained from seed samples. In all cases except Arabidopsis, data sets were generated from plants grown in multiple environments. SNPs were filtered by p-value thresholds specific to each dataset (Arabidopsis p < 0.001, soybean p < 0.05, rice p < 1e-5, sorghum p < 1e-4, and maize p < 0.01) and, when possible, across multiple environments (Table 2). The p-value thresholds varied for each species to reflect differences in statistical power due to different experimental designs. Linkage ranges estimated from genome wide assessments for each population were used to determine locus sizes and tabulate linked genes (Table 1).

**Table 2.** Ionomic loci from GWAS across five species.

| Table 2. Ionomic loci from GWAS across five species. |  |  |  |  |  |
| --- | --- | --- | --- | --- | --- |
|  | Arabidopsis | soybean | rice | sorghum | maize |
| Environments | single | multiple | multiple | multiple | multiple |
| boron (B) | 0 | 401 | 0 | 63 | 10 |
| sodium (Na) | 118 | 331 | 40 | 53 | 0 |
| magnesium (Mg) | 65 | 355 | 10 | 163 | 54 |
| phosphorus (P) | 58 | 372 | 14 | 87 | 39 |
| sulfur (S) | 57 | 362 | 13 | 78 | 54 |
| potassium (K) | 49 | 380 | 12 | 93 | 53 |
| calcium (Ca) | 54 | 360 | 16 | 102 | 43 |
| iron (Fe) | 55 | 354 | 20 | 210 | 67 |
| manganese (Mn) | 59 | 380 | 15 | 175 | 72 |
| cobalt (Co) | 149 | 373 | 0 | 106 | 105 |
| nickel (Ni) | 0 | 374 | 0 | 73 | 56 |
| copper (Cu) | 53 | 353 | 26 | 82 | 72 |
| zinc (Zn) | 51 | 384 | 24 | 103 | 58 |
| arsenic (As) | 68 | 360 | 20 | 0 | 28 |
| selenium (Se) | 126 | 359 | 0 | 114 | 7 |
| rubidium (Rb) | 72 | 352 | 0 | 53 | 59 |
| strontium (Sr) | 60 | 387 | 0 | 0 | 37 |
| molybdenum (Mo) | 109 | 362 | 70 | 115 | 73 |
| cadmium (Cd) | 114 | 381 | 152 | 98 | 11 |
| Total loci | 1404 | 7308 | 432 | 1768 | 905 |

To help identify candidate genes linked to elemental uptake loci in plants, we had previously compiled a list of known ionomic genes (KIG) experimentally shown to alter elemental uptake in plants [26]. As these genes were sourced from the primary literature, we called this our “primary” KIG list. Since the original publication, we have added 19 new primary genes to the list (S1 Table; https://github.com/danforthcenter/KIG_v2), for a total of 201 primary genes. Most primary genes are in Arabidopsis (59%) andrice (33%) with a small number contributed from work in maize, *Medicago truncatula*, and *Triticum aestivum* (wheat). Loci and gene overlaps were considered on a trait-by-trait basis, i.e., Fe uptake GWAS loci were only compared to KIG genes listed for Fe. The overlap between the identified GWAS loci and the KIG identified alleles at genes already known to affect changes in the ionome (Fig 1). Permutations were used to determine the statistical significance of these overlaps by selecting an equal number of random base pair locations across the genome and assigning a “locus range” randomly selected from the actual loci. The primary KIG list covered a low percentage of GWAS loci (<2%) but overlapped with ionomic loci more often than the random locations (80th percentile in Arabidopsis, 95th in rice, >99th in maize).

**Fig 1.**
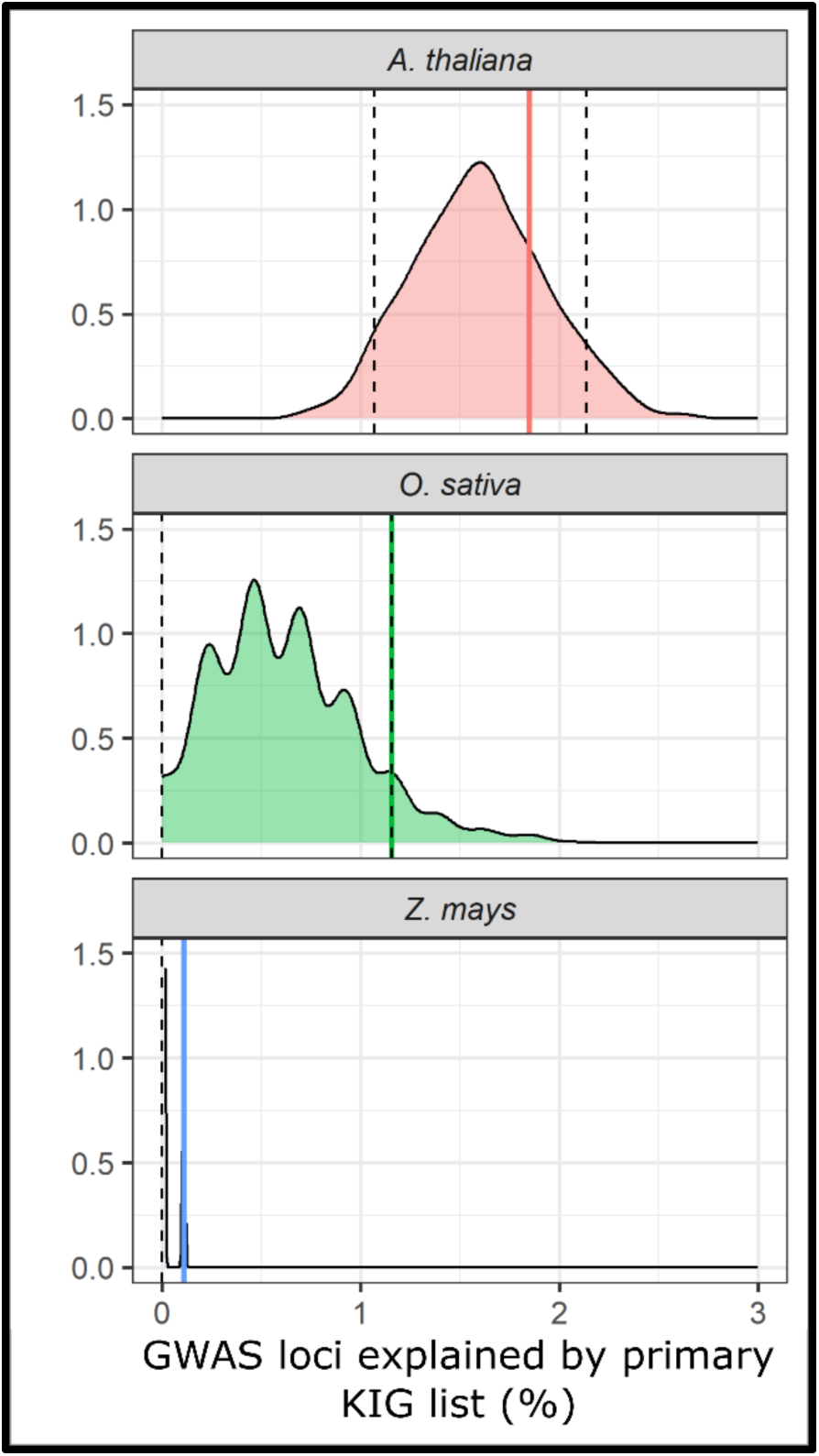
Primary KIG list coverage of ionomic GWAS loci. Solid, vertical lines represent the percentage of GWAS loci explained by a gene from the primary KIG list. Kernel Density Estimate (KDE) plots display the distribution of 1000 random permutations with the 5th/95th percentiles shown as dashed lines. Permutations were made with random genome selections of the same size and species composition as the actual GWAS datasets (S3 Table).

### Can orthology improve the KIG list?

KIGs overlapped the ionomic GWAS loci more often than random genome locations but provided no insight regarding likely candidate genes at 98% of GWAS loci. We hypothesized that inferring orthologs across species could increase the overlap between GWAS loci and expand the list of KIGs and orthologs. Orthologs are genes across species inherited from a common ancestor. Although now in separate species, orthologs tend to retain similar functionality [46], making them good candidate genes when their function is known in one species. Indeed, for many KIGs, orthologs have been tested in multiple species and found to encode conserved functions affecting element accumulation for S, Fe, Cd, Na, K, Rb, Mo, Ni, Cu, P, Zn, As, B, Mn, and aluminum (Al) [26]. For example, alleles at orthologous genes encoding the *MOT1* transporter affect variation in the level of Mo in multiple species [3,28,34,47] and the same has been observed at *HKT1* orthologs for Na [29,48,49]. Inferring the orthologs of KIGs across all of our study species should allow us to discover alleles in genes affecting the ionome, including genes that have not yet been tested in every species in our list.

Ortholog inference algorithms vary by their use of pairwise similarity scores, phylogenetic analysis, and multiple sequence alignment - all of which produce variations in the ortholog groups. We previously published an expanded KIG list [26] made using InParanoid [50,] which uses species-pair comparisons, because the ortholog tables were readily available on Phytozome [45]. For this analysis, we switched to OrthoFinder v2.0 [44] because it performed better than other ortholog group inference methods in benchmarking tests as determined by Emms et al. (2019) and Emms et al. (2020), and forms ortholog groups between 3 or more species. With the ortholog tables from OrthoFinder, we produced “extended” KIG lists that identify orthologs of the primary KIGs in the original species and any species with a sequenced genome, including the five we are studying here. Expanding the list of taxa in the extended KIG list created soybean and sorghum gene lists so we could repeat the KIG/GWAS overlap comparison in all five of our ionomic GWAS species.

The extended KIG lists improved loci coverage across all five species and resulted in significant overlap between the observed GWAS loci and the KIG ortholog lists (>95th percentile) for all species (Fig 2). As expected, this is true whether we randomized the loci (Figs 1 and 2) or randomized the genes in the list (Supplemental Fig 5). While the likelihood that we observed an enrichment of true positives was significant (Fig 2), per-gene-set assessment of false discovery rates (FDR) was relatively high (Table 3), partly due to the large number of genes linked to GWAS loci. Our FDR calculation is a ratio of expected/observed, where the expected average number of extended KIGs linked to random locations from permuted datasets is divided by the observed number of KIGs linked to GWAS loci. Thus, the overlaps between KIG and GWAS loci represent high-value candidates for alleles affecting ionomic variation at conserved genes (Table 3; S6 Table).

**Fig 2.**
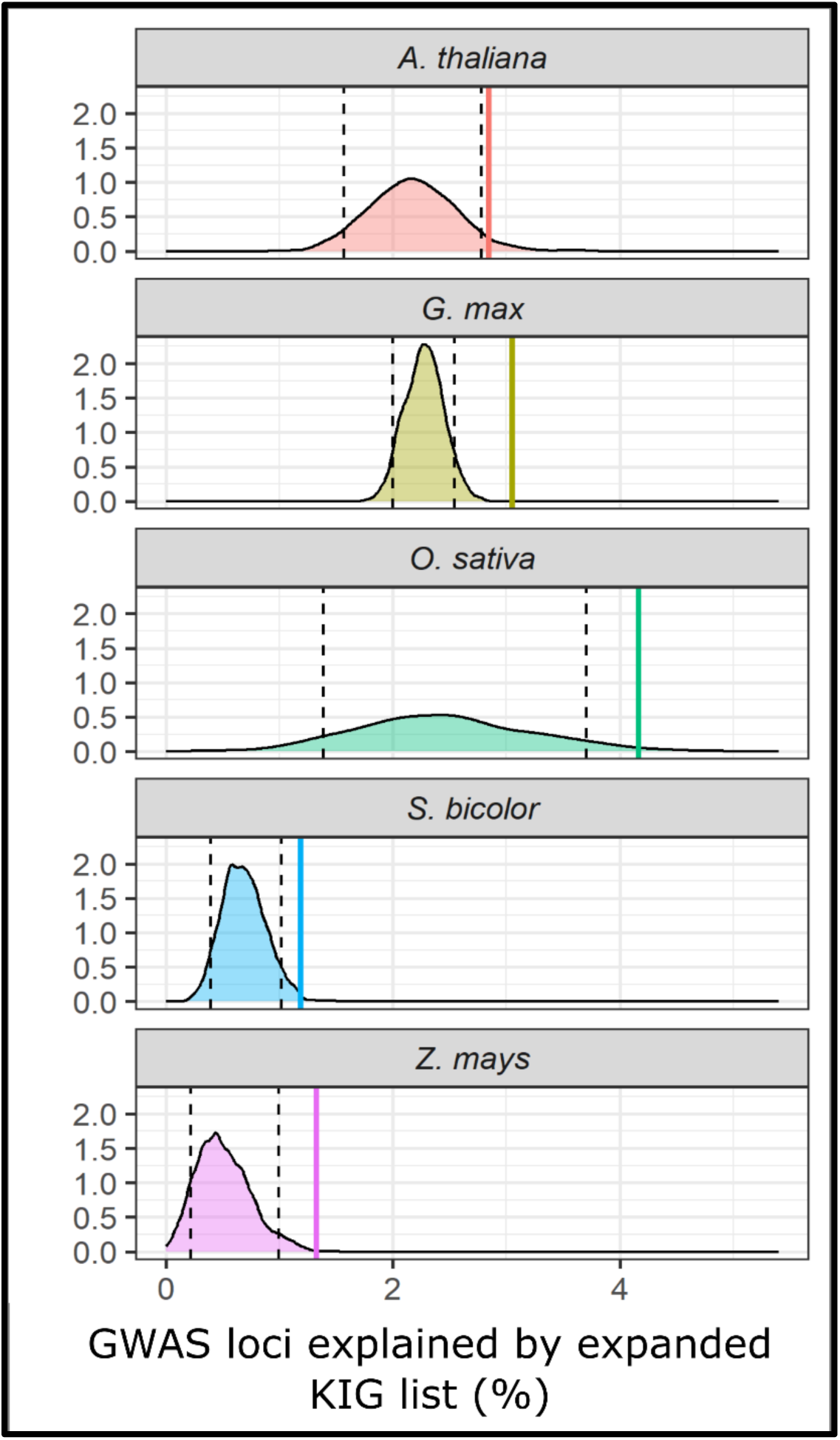
Extended KIG list coverage of ionomic GWAS loci. Solid, vertical lines represent the percentage of GWAS loci explained by a gene from OrthoFinder extended KIG lists. Kernel Density Estimate (KDE) plots display the distribution of 1000 random permutations with the 5th/95^th^ percentiles shown as dashed lines. Permutations were made with random genome selections of the same size and species composition as the actual GWAS datasets (Table 2).

**Table 3.** Summary of extended KIG list overlaps with GWAS loci.

| Species | GWAS loci | random locations | Extended genes | KIG/GWAS overlaps |  | KIG/random overlaps |  | False discovery rate (FDR) |  |
| --- | --- | --- | --- | --- | --- | --- | --- | --- | --- |
|  |  |  |  | genes | loci | genes | loci | genes | loci |
| Arabidopsis | 1404 | 1404 | 163 | 45 | 40 | 32.86 | 30.55 | 0.73 | 0.76 |
| soybean | 7308 | 7308 | 378 | 264 | 223 | 170.03 | 166.23 | 0.64 | 0.75 |
| rice | 432 | 432 | 181 | 22 | 18 | 11.57 | 10.63 | 0.53 | 0.59 |
| sorghum | 1768 | 1768 | 201 | 23 | 21 | 13.36 | 12.13 | 0.58 | 0.58 |
| maize | 905 | 905 | 222 | 13 | 12 | 4.89 | 4.59 | 0.38 | 0.38 |

Among the extended KIGs that overlap GWAS loci are conserved genes known to regulate elemental accumulation across these plant species (Table 4; S4 Table). Identification of alleles linked to orthologous candidate genes via GWAS for the same element provides evidence that these genes not only encode conserved determinants of elemental homeostasis but also indicate that phenotypically relevant, standing variation exists in these genes that could be used to enhance plant mineral nutrition (e.g., P, Fe, Zn, Mn, Mo), improved mineral nutrition for humans (e.g., Fe and Zn), or increased food safety (e.g., Cd).

**Table 4.** KIG ortholog groups overlapping GWAS loci. KIGs were only compared to GWAS loci for traits they were cited for in the primary literature. KIG citations are listed in S1 Table.

| Known ionomic gene family | Cited traits | GWAS trait | Orthologs |
| --- | --- | --- | --- |
| *AtPEN3* | Cd | Cd | *AT3G16340*<br>*Glyma.07G015800*<br>*LOC_Os01g52560, LOC_Os06g36090* |
| *AtHMA2, AtHMA3, AtHMA4, OsHMA3* | Zn<br>Cd | Cd | *AtHMA4, AtHMA2, AtHMA3*<br>*Glyma.09G055600*<br>*OsHMA3*<br>*Sobic.002G083000, Sobic.002G083100*<br>*Zm00001d005189, Zm00001d005190* |
| *AtZIF1, AtZIFL2*<br>*OsZIFL12*<br>*ZmYS3* | Cs<br>Zn<br>Fe | Fe | *AtZIFL2*<br>*Glyma.06G313600, Glyma.06G313700*<br>*Sobic.008G015700* |
| *AtBTS* | Fe<br>Mn<br>Zn | Fe | *Glyma.05G237500, Glyma.08G044700*<br>*Sobic.009G222200*<br>*Zm00001d038916* |
| *AtESB1* | Ca, Mn, Zn,<br>Na, S, K, As,<br>Se, Mo | Mn | *AT1G07730, AT2G39430*<br>*Glyma.08G258300, Glyma.12G067400, Glyma.18G282600*<br>*Sobic.003G064800* |
| *AtNRAMP3* | Fe<br>Mn<br>Zn | Mn | *AtNRAMP3*<br>*Glyma.04G044000*<br>*Sobic.001G170000*<br>*Zm00001d030846* |
| *AtMTP11* | Mn | Mn | *AtMTP11*<br>*Glyma.18G286100*<br>*Sobic.003G349200*<br>*Zm00001d042939* |
| *AtMOT2* | Mo | Mo | *AtMOT2*<br>*Glyma.08G208700*<br>*LOC_Os01g45830*<br>*Sobic.003G237000*<br>*Zm00001d044068* |
| *AtMOT1*<br>*MtMOT1.2, MtMOT1.3*<br>*OsMOT1;1* | Mo | Mo | *AtMOT1*<br>*Glyma.06G074100, Glyma.17G203000, Glyma.17G203200,*<br>*Glyma.17G203500*<br>*Zm00001d033053* |
| *AtPHT1;1, AtPHT1;4,*<br>*AtPHT1;5*<br>*OsPT2* | P<br>As<br>Se | P | *AtPHT1;4*<br>*Glyma.10G186400, Glyma.10G186500*<br>*LOC_Os08g45000* |

The KIG orthologs linked to GWAS loci include several transporters with known roles in element uptake, sequestration, and remobilization. Notably, alleles were discovered in orthologs of transporters in the zinc-induced major facilitator superfamily (*ZIFL1* and *ZIFL2*) genes that affect the accumulation of Fe in seeds (Table 4). Fe is a human nutrient that is limited in many diets [51–53]— making the discovery of these alleles high-priority targets for breeding efforts to improve nutrient accumulation in soybean and sorghum (Table 4). KIGs *MOT1* and *MOT2* encode molybdate transport proteins [28,54–57] and overlap Mo loci in multiple species. Alleles affecting the accumulation of the toxic heavy metal Cd were discovered in all five species at the heavy metal ATPases *AtHMA2*, *AtHMA3*, *AtHMA4*, and *OsHMA3*, which belong to a subset of *HMAs* specific to Zn/Cd transport [11,58–60]. In addition to direct Cd transport, alleles at *PEN3/PDR8*, implicated in Cd transport via chelation with the glutathione-derived phytochelatin [61], affected Cd in three species. The alleles in these transporters and chelator biosynthetic enzymes represent immediate breeding targets for improved human food safety. The phosphate transporter (*PHT1*) family has been shown to affect phosphate and As accumulation in Arabidopsis [62–64] and Se uptake in rice [65]. Orthologs of this transporter were linked to alleles that affect P accumulation in Arabidopsis, soybean, and rice. Thus, these alleles represent possible targets to enhance nutrient uptake for this yield-limiting nutrient. *MTP11* encodes a Mn transporter involved in Mn tolerance and detoxification with the Golgi apparatus [66], and orthologs to *AtMTP11* are linked to Mn loci in four species. Alleles in the orthologs of yet another transporter family, including *AtNRAMP3*, which is involved in Mn homeostasis via the mesophyll vacuoles [67], were also associated with Mn accumulation in Arabidopsis, soybean, sorghum, and maize.

Alleles at regulatory genes from conserved ortholog groups were also identified among multiple species. These included alleles at orthologs of the transcription factor BRUTUS (*BTS*), which affects the expression of genes affecting Fe accumulation in Arabidopsis, including the transporter *IRT1* and the regulatory transcription factor *FIT1* [22,68,69]. Alleles linked to the *BTS* orthologs affected Fe accumulation in soybean, sorghum, and maize (Table 4), demonstrating conservation of function for this transcription factor family as regulators of iron accumulation response across species with substantial variation in Fe transport mechanisms and biochemically distinct Fe chelation systems. As these alleles affect the accumulation of Fe, a nutrient that limits human health, the alleles described here are alleles of immediate value for breeding food staples with improved seed Fe and benefiting global human nutrition. Our approach also detected alleles at a regulator of the ionome, the *Enhanced Suberin 1* (*ESB1*) gene, a dirigent-like protein previously only known via a knockout mutant in Arabidopsis which exhibits altered Casparian strip biosynthesis [70], Ca, Mn, Zn, Na, S, K, As, Se, and Mo levels [71]. GWAS detected alleles at this gene affecting Mn in Arabidopsis, soybean, and sorghum (Table 4), demonstrating that this gene performs conserved functions in regulating ion levels across species and that disruption of the Casparian strip can have similar impacts on the ionome across species with dramatically different root systems and root vascular anatomies.

This analysis demonstrated the utility of combining GWAS across species by using orthology and conservation for genes of known functions. The number of GWAS loci overlapping a KIG ortholog (Fig 2, Table 3) was greater than the overlap with a primary KIG gene (Fig 1, S3 Table). However, this overlap only captured less than 5% of the ionomic GWAS loci. To identify high-quality candidates for the remainder of the GWAS results, we sought a formal procedure for assessing the likelihood of GWAS results at orthologous loci across these five species regardless of previously demonstrated functions.

### Orthology as a key factor for candidate selection

We hypothesized that conservation of orthologs within GWAS loci across multiple species could be a criterion to identify genes of previously unknown function. In the expanded KIG ortholog/GWAS overlap comparison (Fig 2, Tables 3 and 4, S4 Table), we frequently detected phenotypically-impactful alleles in multiple species’ GWAS. GWAS is “forward” genetics and will also detect alleles at genes of previously unknown functions. Therefore, ionomic GWAS datasets will detect alleles at unknown regulators of elemental accumulation. Alleles at conserved but previously unknown regulators of the ionome should also appear more than expected by chance in comparisons between GWAS loci from multiple species.

To test this hypothesis, and identify conserved ionomic genes, we developed a method we call FiReMAGE: Filtering Results of Multi-species, Analogous, GWAS Experiments. This approach identifies the orthologous genes linked to GWAS associations in multiple species and compares their occurrence to a background distribution to assess the false discovery rate. This increases confidence in specific candidate genes linked to GWAS loci without relying on previously known functions. FiReMAGE works by tabulating the genes within trait loci in each species, determining their orthologous relationships across species via OrthoFinder, and retaining ortholog groups with candidate hits from at least three species (Fig 3). A threshold of three or more species was used to reduce the false positives due to the high synteny between some species pairs (e.g.. Sorghum and maize). Like the permutations used with the KIG/GWAS comparisons, FiReMAGE creates 1000 permutations equivalent to the GWAS data by randomly selecting an equal number of genomic locations and ranges from each species’ genome. FiReMAGE runs the permutated sets through the same orthology comparison as the actual data. By comparing the expected (permuted) and observed (GWAS data) orthologs returned from these genomes, this background distribution evaluates the significance of the observed overlaps and provides a false discovery rate. This is a more accurate null distribution for comparison than determining the number orthologous genes in a set of randomly selected genes. Perfectly matched permutation sets were calculated for each set of species and traits allowing FiReMAGE to output FDR for every comparison set (e.g. error control for the 3/5 set of genes associated with Fe in O. sativa from the rice, maize, Arabidopsis comparison). These can be seen as likelihood weights for the candidates in every comparison and are arrived at by performing the same FDR calculation: average number of GWAS-linked orthologs expected by permutations divided by the observed GWAS-linked orthologs in the actual data.

**Fig 3.**
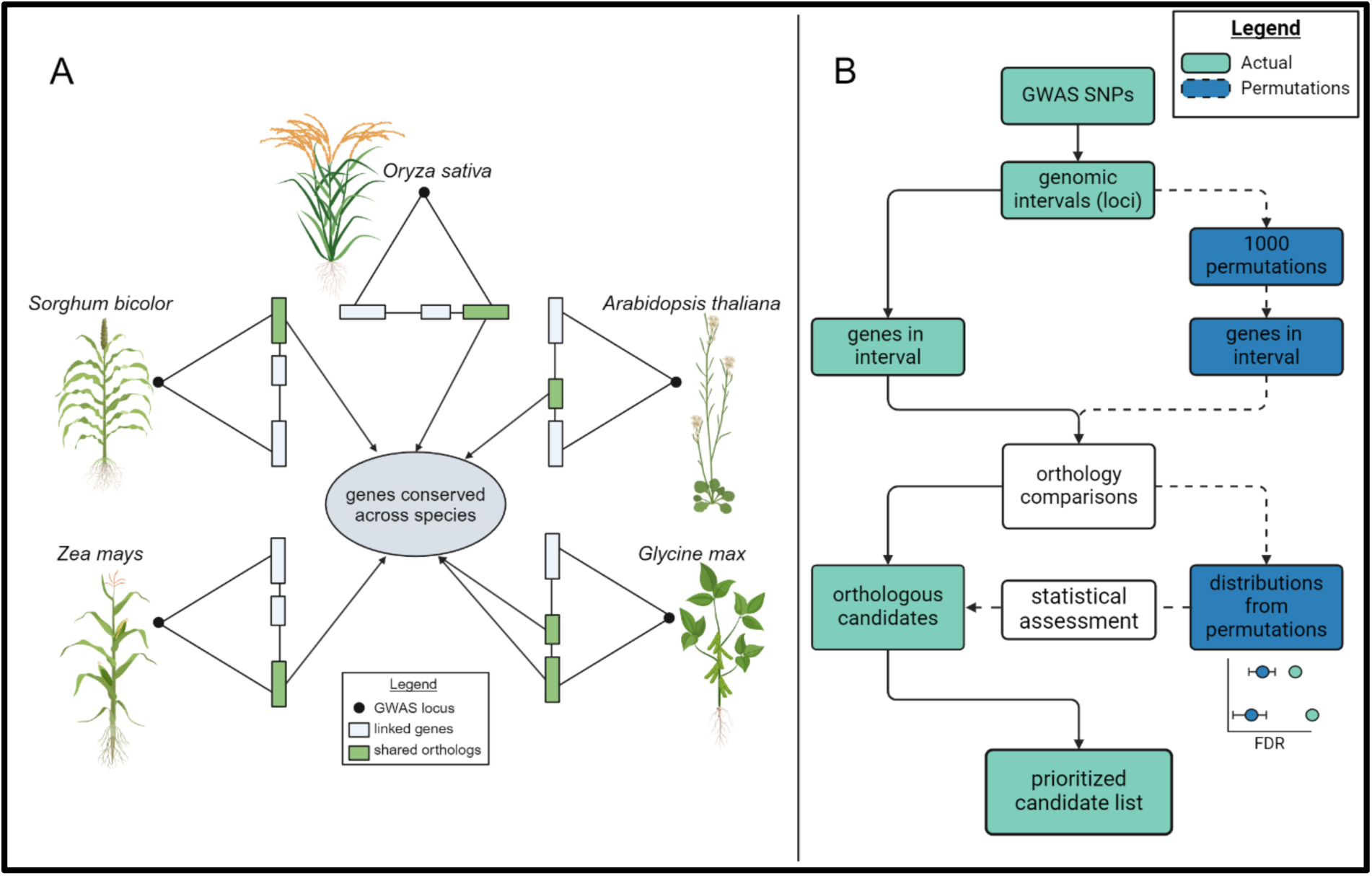
Overview of the FiReMAGE pipeline. (A) FiReMAGE orthology comparison identifies genes linked to SNPs in each GWAS that are also orthologous to each other. (B) FiReMAGE starts by identifying genes within range of GWAS loci intervals. These genes are recorded and filtered through the orthology comparison to identify orthologous candidates. The original GWAS loci then are used to generate 1000 random permutations, which go through the same process as the actual data. Distributions from the permutations are used to assess genes in the real dataset and return a prioritized candidate list. Created in BioRender (https://BioRender.com/c20l148).

With FiReMAGE, we identified candidates in ortholog groups with hits in at least three of the five species for over 30% of soybean and maize loci and over 50% in Arabidopsis, rice, and sorghum (Fig 4). The orthologous loci identified from our GWAS datasets was greater than the 95th percentile of the permutations in each species demonstrating enrichment of conserved genes underlying GWAS hits. However, the individual candidate gene false discovery rates (the ratio of genes expected from permutations/genes observed in the data) was still relatively high (Arabidopsis = 0.58, soybean = 0.43, rice = 0.62, sorghum = 0.39, maize 0.67). Across the 19 elemental traits for the 3 out of 5 species comparison, 3866 unique orthologs were linked to loci in Arabidopsis, 7852 in soybean, 2630 in rice, 2982 in sorghum, and 872 in maize (S5 Fig; S7 Table).

**Fig 4.**
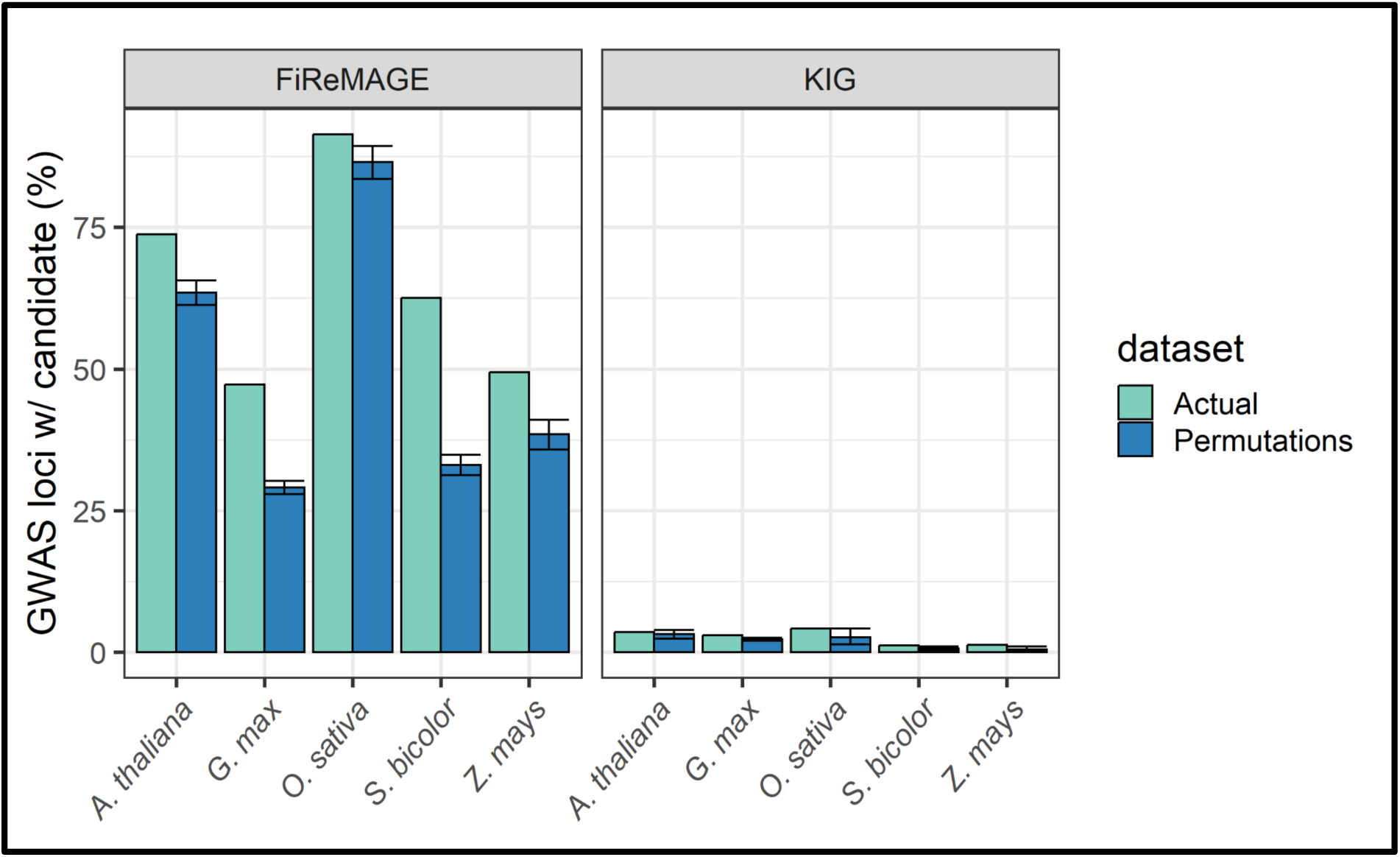
FiReMAGE identifies candidates for a larger portion of GWAS loci than the KIG list. Percentage of ionomic GWAS intervals with a candidate from FiReMAGE’s comparative approach (left) and the OrthoFinder KIG list (right). The actual GWAS dataset is in green; 1000 random permutations of the actual dataset are in blue, with error bars representing 5th/95th percentiles.

The size of candidate lists and false discovery rates vary between species and traits (Figs 5 and 6, S1 and S2 Figs). Requiring more species to be present in an ortholog group returned shorter candidate lists (Fig 5) with lower false discovery rates (Fig 6). While still encompassing a large number of genes, the 3/5 list is a substantial reduction in the number of potential candidates.

**Fig 5.**
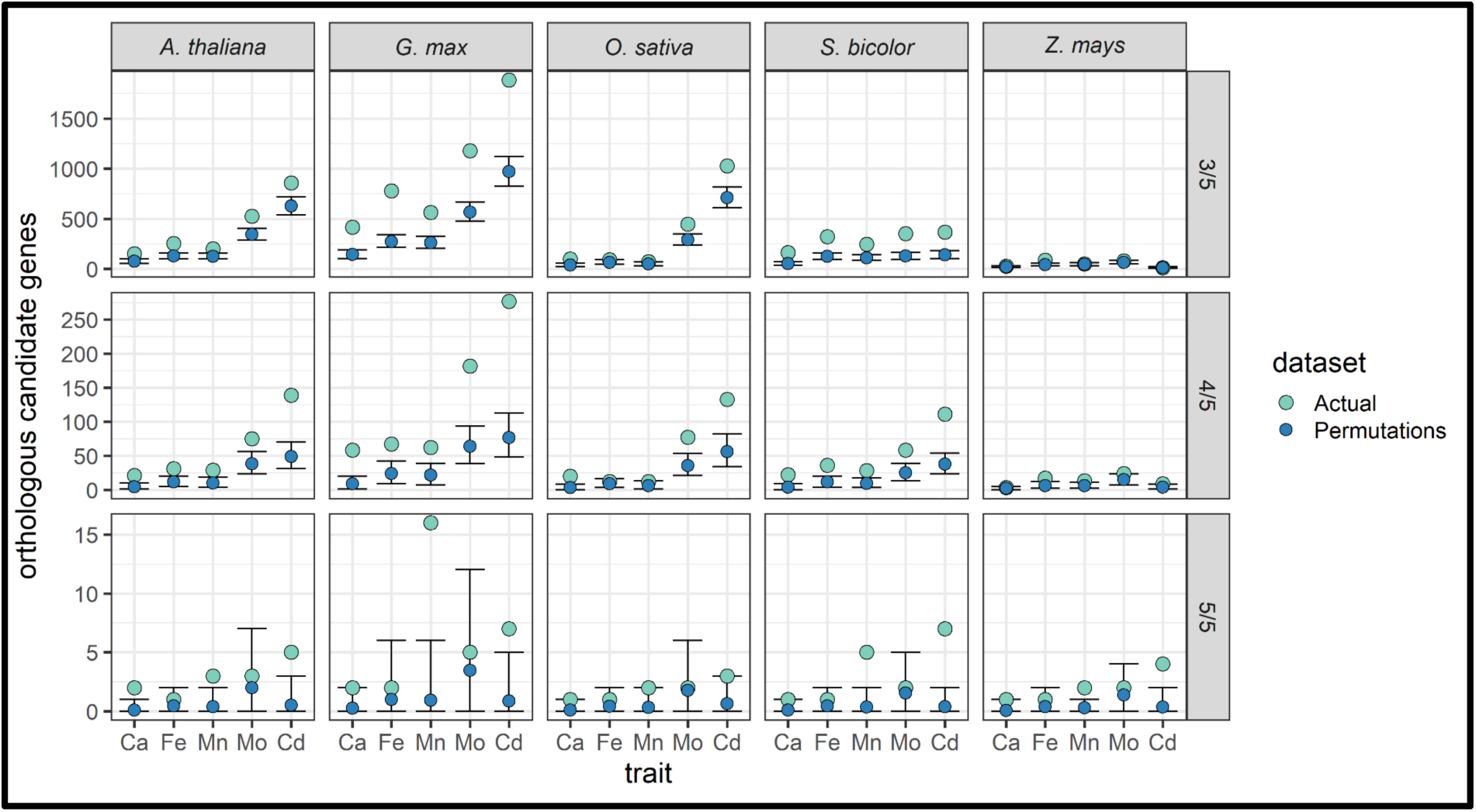
Orthologs from ortholog groups linked to Ca, Cd, Fe, Mn, or Mo loci in 3/5, 4/5, and 5/5 species. Actual dataset counts are in green, and 1000 permutations of the actual data are in blue, with the point representing the mean and error bars representing the 5th/95th percentiles. Columns are by species; rows are the number of species represented in the returned genes’ ortholog groups. Distribution plots for ionomic traits found only in 3/5 or 4/5 species ortholog groups are in S1 and S2 Figs.

**Fig 6.**
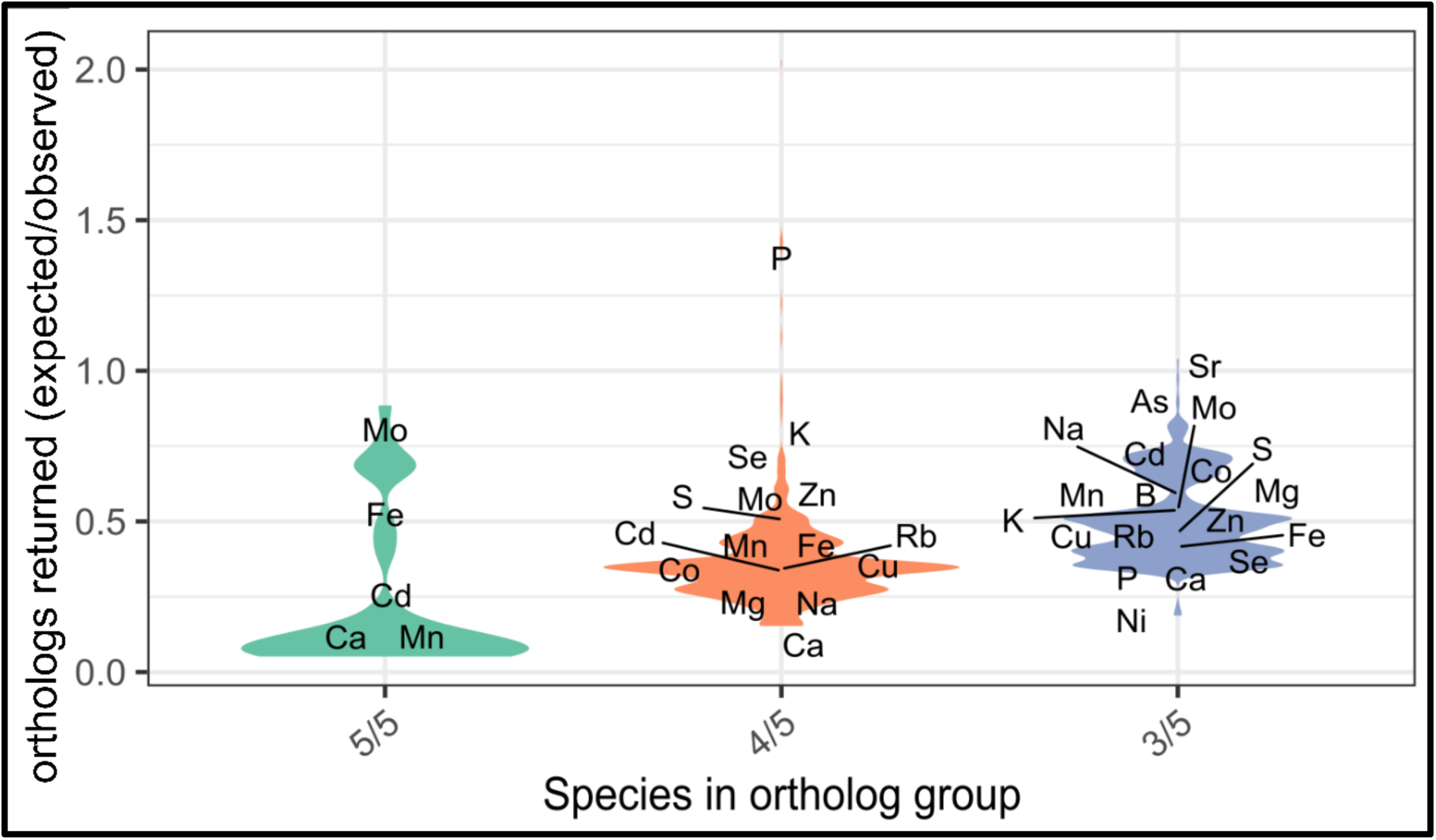
False discovery rate (FDR) of FiReMAGE’s selected candidate genes from ortholog groups with representation from 3/5, 4/5, and all five species for each ionomic trait. FDR was calculated by taking the average number of orthologous candidates returned in the 1000 permutation datasets and dividing it by the number of orthologous candidates returned in the actual dataset for each combination of species, ionomic trait, and species representation in the ortholog group.

High synteny between two species results in many ortholog pairs (syntelogs) at each locus. We carefully examined the influence that the maize-sorghum comparison had on our FiReMAGE results as this was the most closely related species-pair in the data set. We expected the increased synteny between maize and sorghum to increase the number of orthologous genes found (highlighted red in Fig 7). This might increase the number of false positive (FDR) ortholog/GWAS overlaps and obscure the signal at true causative genes. This high synteny did not, however, dominate the results. GWAS overlap at ortholog groups from species with increased synteny do not make up the majority of the loci detected. The largest intersection with maize, sorghum, and third species (maize-sorghum-soybean) detected 245 ortholog groups, and was only the fifth largest intersection out of all ten ortholog group species combinations. Additionally, the FDR of these genes is comparable to 3/5 ortholog groups derived from species with less synteny, suggesting that this is not a major concern for this experiment.

**Fig 7.**
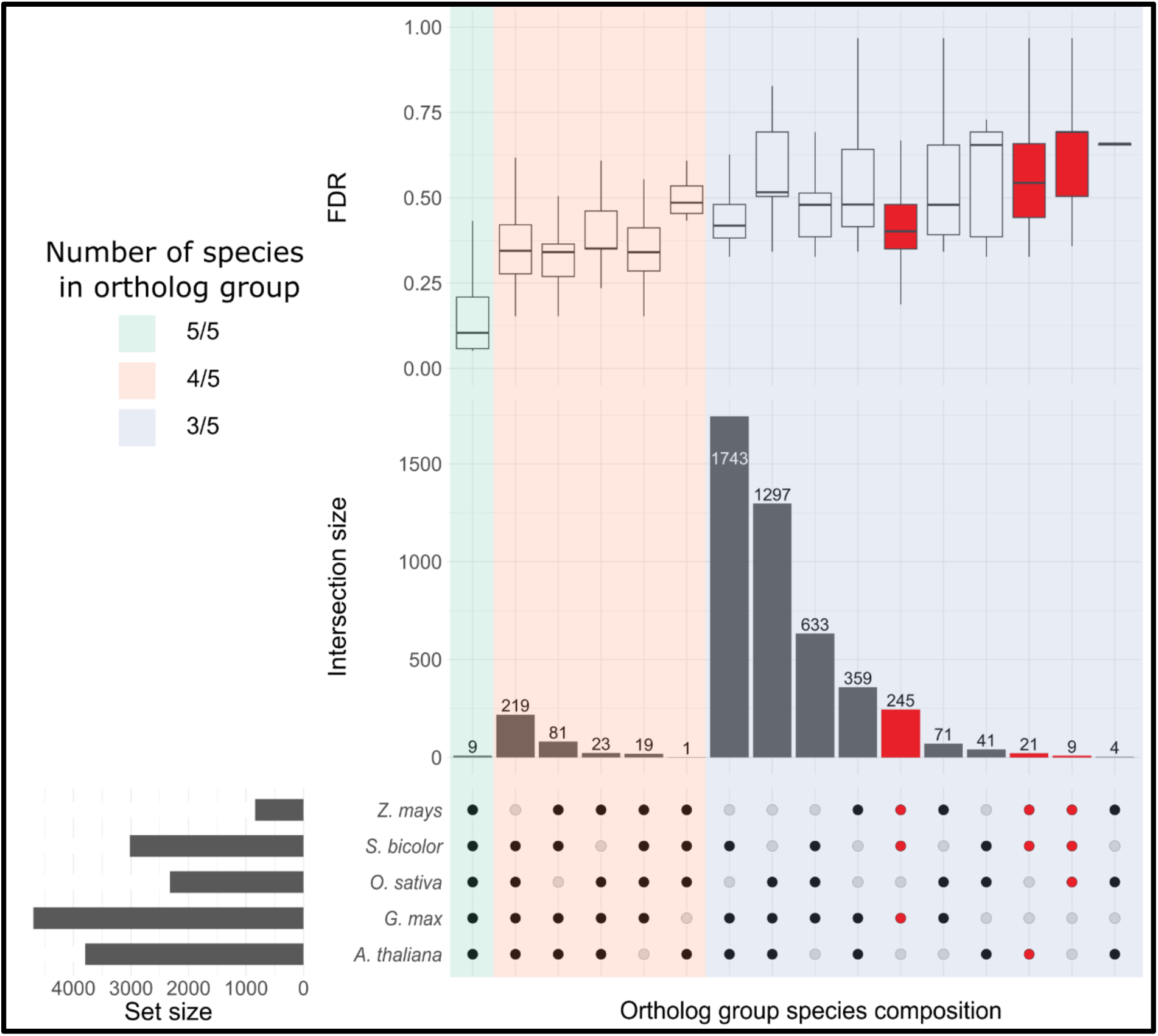
UpSet plot of FiReMAGE’s selected candidates’ ortholog group composition. The intersection matrix shows the unique compositions of all ortholog groups determined by OrthoFinder. Ortholog groups are ordered by the degree (number of species) of the intersection and then by group size. Intersection size represents the number of ortholog groups matching the species composition below. Set size shows the total number of instances in which a species appears in all ortholog groups. The top panel displays the individual genes’ false discovery rate (FDR) distribution in their respective ortholog groups for all ionomic traits. Note that some genes may be represented twice in the boxplot distribution if they appear for multiple traits. Thus, they may have multiple FDRs due to the different SNP distributions and FDR of the ionomic traits. Because of the theory that high levels of synteny between sorghum and maize may increase the number of false positives returned, ortholog groups with two of the three species being sorghum and maize are highlighted red.

### FiReMAGE identifies highly likely candidates and novel hypotheses for the regulation of element accumulation

For each ionomic trait and species combination, FiReMAGE produced a list of orthologous candidates and an FDR estimated by permutation (S7 Table). If elemental accumulation is controlled by conserved genes, then the FiReMAGE procedure should identify more orthologous gene/loci overlaps than expected by chance via the permutations. FDRs differed across the element accumulation traits and species combinations. Generally, comparisons with low FDRs were observed for elements with a previously demonstrated high heritability [72–76] and with GWAS hits in all five species: Cd, Ca, Fe, Mn, and Mo (Fig 5). Of 23,117 FiReMAGE candidates, only 81 belong to these 5/5 ortholog groups. If we rank all candidate genes using the FDR for each gene set from the perfectly matched permutation scores, candidates 1-56 and 115-116 are linked to GWAS loci at orthologous candidates in all five species. The top 10% of candidate genes ranked in this way correspond to 233 genes from 18 element/species/comparison combinations (S8 Table). That a lower FDR was observed when genes were present in more species demonstrates the value of using orthology, the power of replication across the species, and the utility of our permutation-calculated FDR to rank candidates.

Among the candidates from orthologous groups linked to loci in all five species are some of the most significant GWAS SNPs in the input set. The GWAS loci linked to these 5/5 candidate genes (n = 64) were enriched for the top 10% most significant SNP-trait associations from their respective GWAS (Chi-square p-value > 1.25 e-10; obs: 21 True, 43 False; exp: 6 True, 58 False). Low p value loci within an individual species’ GWAS (n=21), might have been considered targets for candidate verification. FiReMAGE improves upon this by identifying likely causative conserved genes at each of these locations, accelerating the process of gene function discovery. In addition, FiReMAGE also outputs the likely alleles in the species where the effects at these conserved genes were less strong. Thus, by using orthology, FiReMAGE identified genes linked to GWAS SNPs with “middling” signal to identify the remaining conserved candidates.

The identities of the genes encoded by the FIReMAGE candidates in our 5/5 species lists implicate a wide variety of biological processes in elemental homeostasis. The 81 candidates with alleles in all five species belong to nine orthologous groups. For Fe, among the 5/5 candidates with no previously known role in Fe homeostasis, is an ortholog group that encode Mevalonate kinases across these five species, suggesting a currently unknown role for the mevalonic acid pathway in regulating Fe homeostasis in flowering plants. Among the top Cd candidates, we find the *HMA* ortholog group mentioned above in the KIG list (Table 4), RAB GTPase homologs of *RABA1i* that are involved in regulating cell wall composition [77], and an ortholog group of Domains of Unknown Function 567 (DUF567) protein family. In Arabidopsis, DUF567 is expressed in the roots and identified among a list of 129 genes associated with metal hyperaccumulation in *Arabidopsis halleri* [78]. Thus, FireMAGE provides additional experimental evidence, leveraged across five species, that this DUF567 family affects ion homeostasis and may underlie ecological adaptation via hyperaccumulation in Arabidopsis relatives. For Mo, the 5/5 candidates are the molybdate transporters, *MOT2*, listed in the KIG list for Mo accumulation [54], and adenine nucleotide transporters (*NTT1* and *NTT2*).

Nucleotide metabolism is tied to Mo via molybdopterin, which is synthesized from nucleotide precursors [79] to form reaction cofactors for four plant enzymes [80] suggesting a mechanism for these associations. For Ca, the orthologous candidates are annotated as encoding a Ring/Ubox superfamily with no known role in Ca accumulation. Lastly, for Mn, the candidate ortholog groups are annotated as histone proteins and cellulose synthase A9 (*CESA9*). *CESA9* has been shown to influence the cellulose content and cell wall biosynthesis in the epidermal seed coat in Arabidopsis seeds [81]. In cellulose synthesis, Mn ions are used to catalyze glycosidic bond formation and CESA protein coordinates Mn ions with the glycosyltransferase substrate uridine diphosphate glucose [82]. Given the large number of Mn-dependent processes in the cell, the association is unlikely the result of alleles affecting Mn levels via the size of the cellulose synthase sink. It may be that Mn homeostasis in the golgi and trans-golgi networks [83], are sensitive to the binding state for coordinated Mn in CESA. Thus, highly likely candidates identified as conserved genes underlying GWAS peaks in this species set have reasonable proposed mechanisms of action, are worthy of further scientific investigation and, again, represent potential immediate-use targets for breeding and crop improvement.

We also compared the FiReMAGE candidates to another candidate selection method performed in maize: Camoco [17]. From the 47,000 candidate observations derived from multiple expression network and elemental trait combinations, we selected the top 200 observations as ranked by the FDR calculations in Camoco’s multiple expression-network-based prioritizations and ionomics trait analysis [17]. This selection contained 122 unique candidate genes to compare against FiReMAGE’s evolutionary-based prioritizations. Only two of these112 candidates were detected as conserved regulators by FiReMAGE. One gene, *Zm00001d04461*, identified through both methods as affecting Mo, encodes an F-box domain containing protein; the other, *Zm00001d012274*, affecting Fe accumulation, is an enzyme encoding an Omega-hydroxypalmitate O-feruloyl transferase involved in the production of suberinized hydrophobic barriers [84] that are known to influence ionomic traits. Despite the clear mechanism implicated, the finding that only two genes affected variation detected by the two methods indicates that single-species expression networks and orthology selection methods boost different genetic signals to tie GWAS results to candidate genes.

## Discussion

We have demonstrated that combining orthology and GWAS from multiple species detects conserved regulators of phenotypic variation. Using the Known Ionomic Genes (KIG) as a validation set of curated annotations confirmed our hypothesis that combining GWAS and orthology would correctly identify variation affecting elemental accumulation. This resulted in the discovery of phenotypically-impactful alleles affecting ionomic variation in genes of previously known functions encoded in these five species. Additionally, the FiReMAGE approach went beyond the KIG list by identifying numerous candidates in genes without previously determined roles in element accumulation (S7 Table). This ability to identify and prioritize candidate genes for conserved roles in trait variation using the natural variation of multiple species transcends one of the key limitations of the post-genomic era: ascertainment bias due to prior study. Prior research bias has resulted in gene discovery focused on a handful of elements due to researcher interests, leaving a majority of elements under-studied [26]. Similar biases for prior demonstration of gene function, result in the preferential publication of additional alleles at homologs of known function rather than the exploration of natural alleles at locations with no genes of prior molecularly demonstrated effects on the studied trait. The overlap of less than 5% between GWAS outputs and the KIG indicate that the KIG list is vastly incomplete [8], and many more plant genes that regulate the ionome are waiting to be discovered. The overlap between the FiReMAGE output and genes of previously known function as well as the identification of orthologous high likelihood produced a comprehensive public list of alleles at candidate conserved regulators of 19 elemental accumulation for future molecular mechanistic studies or marker-assisted selection (S2 and S7 Tables).

Instead of selecting candidates with prior annotations, FiReMAGE annotates orthologous genes as high-value candidate genes for further molecular function characterization. Experimental error, high genome-wide statistical thresholds, and low statistical power in a GWAS results in high false negative rates (e.g. insignificant p-values at a true causal gene). Meanwhile, every gene linked to SNPs with a significant p-value will not be a true causal gene and assuming so would increase false positives and result in spurious conclusions. The permutation results in Figs 1 and 2 demonstrate the power of collectively assessing GWAS datasets across species. When the same gene is linked to GWAS SNPs in multiple species, these independent experiments provide multiple lines of evidence for that gene’s function and increase the likelihood that it is causal. T

### Orthology-based FiReMAGE is highly effective for identifying gene candidates

The integration of orthogroups into gene annotation provided improved estimates of gene function, as demonstrated by the improved GWAS/KIG overlap when orthologs were included (Figs 1 and 2, Table 3, S3 Table). As a result of this success, we hypothesized that orthology, even in the absence of gene function annotations, would also identify conserved genes as candidates without any prior demonstration of ionomic function. FiReMAGE identified more orthologous genes linked to GWAS loci for the same trait than expected by chance (Figs 4 and 5), demonstrating that this approach works. In addition to KIGs (Table 4), FiReMAGE also returned non-KIG ortholog groups that encoded natural variation in elemental accumulation (S7 Table). These FiReMAGE candidates are likely to encode conserved regulators of elemental accumulation, for which our analysis is the first genetic demonstration of this role. Finding conserved critical regulators of ion homeostasis by GWAS validates the expectation that conserved regulators (e.g., hub genes) will repeatedly accumulate phenotypically effective variation. It also dramatically broadens the capacity of GWAS to work as a gene function discovery tool in the forward-genetics mode. Thus, comparing element accumulation association studies across organisms can help us discover additional element homeostasis mechanisms. Our approach relies on both annotated genomes and GWAS data, but it should be generalizable to any homologous traits across taxa. FiReMAGE does not require any information other than a list of GWAS SNPs and gene positions in genomes, thus avoiding prior knowledge bias and enabling accelerated gene function discovery for traits and species with little to no functional annotations or prior work.

One of the non-KIG ortholog groups with alleles in all five species detected by FiReMAGE for Cd encodes a domain of unknown function 567 protein (DUF567) family. This DUF567 is just one of the 198 elemental accumulation candidate genes identified by FiReMAGE in Arabidopsis that has no previously known function in any biological processes (S9 Table). No other genes whose best annotation is as domain of unknown function (DUF) containing gene are in 5/5 ortholog groups, but 19 are in the top 10% of Arabidopsis candidates for all traits, and 46 unknown genes are in the top 25% when sorted by their permutation estimated FDRs. None of these unknown genes would be considered candidates if relying only on gene annotations. That these DUFs are within the gene sets with the lowest permutation-estimated FDRs which also include known conserved regulators from multiple species (Table 4) suggest these DUFs are conserved, previously unknown, regulators of the ionome. Future single-gene mechanistic experiments may result in the annotation of these genes as elemental regulators. Problems with annotating genes and Gene Ontology (GO) terms in general, and plants in particular, have been extensively discussed [7–9]. FiReMAGE side-steps this problem, as is illustrated by the observation that it finds 602 orthologous gene candidates for Mo and 116 for Rb (S7 Table), yet there are no GO terms for the transport, accumulation, or metabolism of either Mo or Rb.

Using data from multiple experiments, FiReMAGE emphasizes loci that would be passed over in a single-species experiment. In many cases, FiReMAGE outputs strong signals when orthologs are considered across taxa for loci with moderate effects in each species. Five traits, Ca, Fe, Mn, Mo, and Cd (Fig 5, S7 Table), have ortholog groups with candidate gene hits in all five species. The federation of data across multiple species implemented by FiReMAGE can boost the signal at genes encoding alleles that were below the p-value thresholds typically employed for single-species GWAS assessment. This is illustrated by the results located within GWAS loci at the ortholog groups containing *MOT2* and the Cd/Zn transporting *HMAs* (Table 4). Some of the p-values in single species comparisons are very significant, but not all. The worst performing taxa for these ortholog groups were soybean for Mo accumulation at *MOT2* (p-value = 0.0104) and Arabidopsis for Cd accumulation at the tandem gene duplication comprising AtHMA2, AtHMA3, and AtHMA4 (p-value = 5.6e-05). Neither of these associations would meet a multiple testing corrected p-value cutoff for significance if considered alone in their respective single species GWAS (S7 Table). Finding associations at these loci in all five of the species demonstrates that variation at these loci is present. Indeed, AtHMA3 is at a major effect QTL affecting the level of Cd in leaves at a p-value < 10e-20 [11]. The cause of this association, and our detection of a smaller effect on seed Cd accumulation, is a complicated collection of loss-of-function alleles of various frequencies encoded at this locus [11]. One null mutant allele in HMA3 is present in the reference Col-0 accession [58]. This loss of function allele induces sensitivity to metals, including Cd which it directs to the vacuole [85]. The dramatic effect of these alleles on Cd accumulation in leaves and weaker effects in seeds likely results from a strong effect of HMA3 on vacuolar sequestration, and a weaker contribution to Cd remobilization during seed filling in Arabidopsis.

For those genes that are in the KIG orthologs set, these have been highlighted as genes affecting the ionone independently via FiReMAGE, demonstrating phenotypically affective variation co-segregating with identified SNPs, and via prior work on molecular genetic demonstration. These alleles all represent utility-ready breeding targets that are the highest confidence hits in our study. Of particular note are the alleles identified by FiReMAGE that overlap the KIG and alter the levels of minerals known to limit human health. Because the GWAS data sets employed here are all studies of seeds, all of the SNPs affecting variation in elements important for human nutrition can be incorporated into breeding programs. For example, all Fe and Zn alleles at KIG can be put to use immediately in public breeding seeking to increase the concentration of these critical human nutrients to overcome health-limiting deficiencies. Similarly, the Cd and As associations identified in these materials at KIG orthologs (Table S7) can be put to use immediately to reduce the levels of these toxic elements in the food supply.

### FiReMAGE approach and utility for gene discovery

Our FiReMAGE approach combines a wide array of information. It has many steps (functions) and dozens to hundreds of user-defined parameters. Many of these lack obvious correct or best choices for parameter values. These include the p-value thresholds for the GWAS input data sets, the linkage range for SNP-to-gene relationships, and the outgroups for constructing ortholog relationships. In addition, several variables are inherent in the data and experimentation that also affect FiReMAGE, including differences in measurement error in each experiment, variation in the statistical power of each GWAS panel, the total genetic variation in a species, and the association panel selection. By using permutations instead of an idealized null distribution, FiReMAGE estimated how likely the results were to happen by random chance, given this set of input parameters, and provided estimates of the likelihood that our predictions are true. The FiReMAGE approach calculates these permutations for every trait, species, and dataset combination. As researchers get better estimates of parameters or add additional data, the permutations must also be updated. Preliminary testing determined that random selection parameters were replicating real data structure and composition, and our permutations serve as consistent internal controls for benchmarking results.

One of the reasons to employ permutations of regions instead of random sets of genes, was the collinearity of genes in genomes. We expected that the high synteny between the three Poaceae species, rice, sorghum, and maize, would inflate false positives. To the extent that multiple linked genes are shared across taxa due to conservation of gene order, we expected the ortholog groups detected as 3/5 hits with FiReMAGE in only the Poaceae to have high FDR. However, these ortholog groups did not have dramatically increased FDR compared to other 3/5 ortholog groups (Fig 7). The combination of the recent genome duplications and uneven loss of duplicate genes well demonstrated in maize, Arabidopsis, and others [86–91] and the prevalence of chromosomal rearrangements and transposition of genes in plant genomes may have shuffled these genomes enough to permit the detection of single conserved genes within our selected window sizes. Although we did not see a huge influence of synteny in our comparison, other species combinations may find controlling for synteny critical [92]. Any undue influence of synteny will be reflected in the permutation FDR calculations, highlighting the value of empirically estimated expectations over naive null hypotheses.

The success of FiReMAGE with these particular data sets and user-set parameters demonstrates that this approach is viable. One feature of this analysis was the reliance on user-reported p-values for a number of the data sets. This resulted in uneven p-value thresholds and presumably influenced false positive and negative rates due to differences in statistical power and error across the input GWAS experiments. Because FiReMAGE federates data together, a search for the optimal p-value thresholds for each experiment would require determining how changing the input data affected the performance of the comparison by FiReMAGE. One interesting feature of FiReMAGE is that it is highly sensitive to false-negative rates. Using too stringent of a p-value cutoff for each species GWAS results will fail to input into FiReMAGE the true associations of modest effect near conserved regulators. FiReMAGE cannot then include this gene in an overlap and loses one true overlap as a result. This problem increases as the number of species compared goes up, when stringent pvalues are used. Thus, we chose p-value thresholds for the single-species inputs that are considered permissive to reasonably limit false negative rates and still retain signal in this FiReMAGE comparison. Again, the use of permutations to benchmark the comparison and determine significance of FiReMAGE validated this choice. As candidate genes suggested by FiReMAGE are tested, the genes that are confirmed and genes that were false positives will provide an additional mechanism to compare and refine parameters.

The high FDRs from the KIG comparison demonstrate why common practices filter SNPs with stringent thresholds (Bonferroni, FDR, etc.) before referencing functional annotations of linked genes. However, these recommendations do not address that even with generous genome coverage, KIG candidates cannot explain a large portion of ionomic variation (Figs 1 and 2). While some loci highlighted by GWAS at these thresholds are due to spurious associations and some KIGs do not drive elemental phenotypes in all populations in all species, we propose that the majority of loci lack KIG candidates because there are a vast number of gene functions and allele consequences yet to be discovered [8]. Candidate selection methods reliant on prior information have inherently narrow scopes that increase false negatives and, by focusing efforts on highly likely confirmation experiments at genes of known function, inhibit gene discovery. The KIG list is slightly better at discovery, as it is valuable in transferring information between species with high quality annotations to species with fewer genomic/genetic resources. However, identifying novel ionomic candidates or functional annotations requires novel approaches, such as FiReMAGE.

### Limitations of FiReMAGE

One effect of the FiReMAGE approach on candidate gene assessment is that it ignores alleles and false positives that occur in GWAS results from a single species or in a limited number of species. While of great potential utility, this approach can potentially contribute to false negative assessments in at least two ways. Firstly, the FiReMAGE approach will necessarily miss taxonomically-restricted solutions and limitations to element homeostasis. For example, glucosinolates in Arabidopsis, benzoxazinoids in maize, and nodulation in soybeans, all have impacts on the ionome [93,94] and are not evolutionarily conserved processes across these taxa. As a result, alleles affecting variation in these processes, even if they have substantial impacts in one taxon, would not be prioritized by FiReMAGE. The KIG list or more involved experiments like gene transcript expression would be better selection methods in these cases [17]. Secondly, the requirement to appear in multiple lists and the stringency of the criteria for inclusion in the GWAS outputs contribute to false negatives intrinsic in FiReMAGE. Researchers particularly worried about false positive assessments in their GWAS outputs may not have considered the effect that the high rate of false negatives will have on downstream meta-analysis and data federation approaches, like FiReMAGE. Because detecting conserved candidates by FiReMAGE relies on true positives in multiple species, providing too-stringent lists of loci will result in poor performance as more species are added to the comparison. Communities that make raw data readily available for re-analysis are likely to see approaches like FiReMAGE return better results.

Another limitation is that FiReMAGE requires strict trait matches across taxa and we only compared KIGs to loci of traits for which they had been cited. Many elemental uptake traits are related and often elements are co-transported by the same transporters. The same gene could have different elemental phenotypes in taxa due to many factors, including the interaction with species-specific mechanisms (see above). The rules governing elemental interactions are poorly understood [95,96]. Rather than attempting to relate ortholog groups to similar elements they have yet to be tested for with our (likely) biased understanding of the ionome, we chose a strict comparison between traits across species. We have provided a list of KIGs overlapping any trait loci (S4 Table) to guide future researchers interested in exploring their potential functions in related traits.

A functional requirement for FiReMAGE is the collection of the same trait across multiple diverse taxa. Many traits, such as flower morphology and resistance to specific pathogens, are not comparable across species. Additionally, collecting trait data from well-powered (i.e. large) GWAS populations across multiple species can be time and resource-intensive. However, advances in phenotyping technologies are enabling the creation of multi-species datasets, and more datasets are publicly available in GWAS databases like AraGWAS [97] for Arabidopsis, RiceVarMap2 [98] for rice, and GWAS Atlas [99] for studies in cotton, maize, soybean, rice, and sorghum. FiReMAGE is able to identify candidates for any trait where the data is available in three or more species. We note that the threshold of three or more species excludes eudicot-specific candidates in this five-species comparison. As more eudicot species with ionomic GWAS experiments become available it may be possible to carry out FiReMAGE searches for candidate genes using phylogenetically targeted subsets of taxa.

Lastly, because FiReMAGE relies on orthology, the limitations surrounding current ortholog group inference methods apply to the candidate lists produced by FiReMAGE. While OrthoFinder v2.0 is the best algorithm for whole genome comparisons [44], there are likely some gene trees with evolutionary histories that make accurate ortholog inference challenging [100]. We recommend following best practices for operating OrthoFinder (see methods), including proper species sampling and validating gene tree phylogenies for FiReMAGE candidates of interest when appropriate.

### Future mutant screens

Unbiased candidate selection methods are necessary to transition into research using non-model species and with genes of unknown function. Our approach has identified the conserved genes associated with genetic variation via analogous GWAS in multiple species and their likelihood as causal genes in ionomic phenotypic variation. In future studies, testing mutant knock-out libraries in Arabidopsis [101], sorghum [102], and maize [103] will allow us to validate FiReMAGE predictions and improve candidate selection parameters.

Our research demonstrates that the disparate approaches of the KIG list and FiReMAGE both present valuable ionomic candidates, each with distinct advantages and weaknesses, and highlight the importance of strategic decisions surrounding candidate gene identification for downstream quantitative genetic analysis. Future experiments will further explore best practices and outcomes of comparative GWAS methods, including parameters to balance false negatives and false positives. We also can imagine implementing a ranking equation that includes variables other than permutation calculated false discovery rates. We hope this tool facilitates gene discovery in genetics experiments across all organisms.

## Acknowledgements

We thank Ryan Hartsock for contributions to the early development of our orthology comparison methods.

## Supplemental Files

**S1 Fig. Orthologs in groups with 3/5 species representation.** Error bars represent the 5th/95th percentiles of the permutations.

**S2 Fig. Orthologs in groups with 4/5 species representation.** Error bars represent the 5th/95th percentiles of the permutations.

**S3 Fig. Distribution of list sizes in the inferred KIG permutations depending on ortholog tables.** The old distribution of permutations’ list sizes was from using OrthoFinder to call ortholog groups only between the species in the inferred KIG. The new distribution of permutations’ list sizes shows the list size when *Liriodendron tulipifera* is added to our OrthoFinder run as an outgroup to our target species. The green dot represents the number of inferred KIGs in the actual list.

**S4 Fig. Inferred KIG and random permutations of the KIG list compared to GWAS loci.** Solid, vertical lines represent the percentage of GWAS loci explained by a gene from OrthoFinder inferred KIG lists. Kernel Density Estimate (KDE) plots display the distribution of 1000 random permutations with the 5th/95th percentiles shown as dashed lines. Inferred KIG list permutations were made from the orthologs of a list of random genes equal to the species and elemental composition of the primary KIG list (see Random permutations methods section).

**S5 Fig. Distribution of the number of orthologous candidates returned per loci.**

**S1 Table. The current primary KIG list.** Genes with functional characterizations from primary literature sources that are linked to changes in the ionome.

**S2 Table. Updated KIG list with OrthoFinder inferred orthologs.** *T. aestivum* (wheat) is not included in this version of the inferred KIG list.

**S3 Table. False discovery rates of the genes and loci overlaps in Fig 1**. False discovery rates are calculated by taking the average number of genes/loci in random permutation overlaps and dividing by the number of genes/loci in the actual GWAS datasets.

**S4 Table. KIGs overlapping GWAS loci.** KIGs compared to all elemental uptake trait loci regardless of KIG list citations.

**S5 Table. KIGs in FiReMAGE’s candidate lists.**

**S6 Table. False discovery rate table for gene and loci overlaps in S4 Fig.** False discovery rates are calculated by taking the average number of genes/loci in random permutation overlaps and dividing by the number of genes/loci in the actual GWAS datasets.

**S7 Table. The complete FiReMAGE priority lists for each trait and species.**

**S8 Table. Top 10% of all FiReMAGE candidates.** FiReMAGE candidates were sorted by their permutation calculated false discovery rates, and taking the top 10% with the lowest FDR.

**S9 Table. FiReMAGE candidates in Arabidopsis with no previously known biological processes function.** We used ThaleMine [104] to analyze Arabidopsis candidates for each trait to identify genes with unknown function.

**S10 Table. Soybean SNPs associated with ionomic traits.** SNP associations generated with methods outlined in “Soybean GWAS and 2015-2017 field experiments”.

## Notes

https://github.com/danforthcenter/KIG_v2/settings

https://github.com/danforthcenter/FiReMAGE

### Competing Interest Statement

The authors have declared no competing interest.

### Summary of Updates

The manuscript has been rewritten for clarity, adding further details to the methods. No new data or results have been added since the previous version was submitted to BIORXIV,

