## Supplementary figures and images for "A comparative approach for selecting orthologous candidate genes in genome-wide association studies across multiple species"

### S1_Fig.tiff

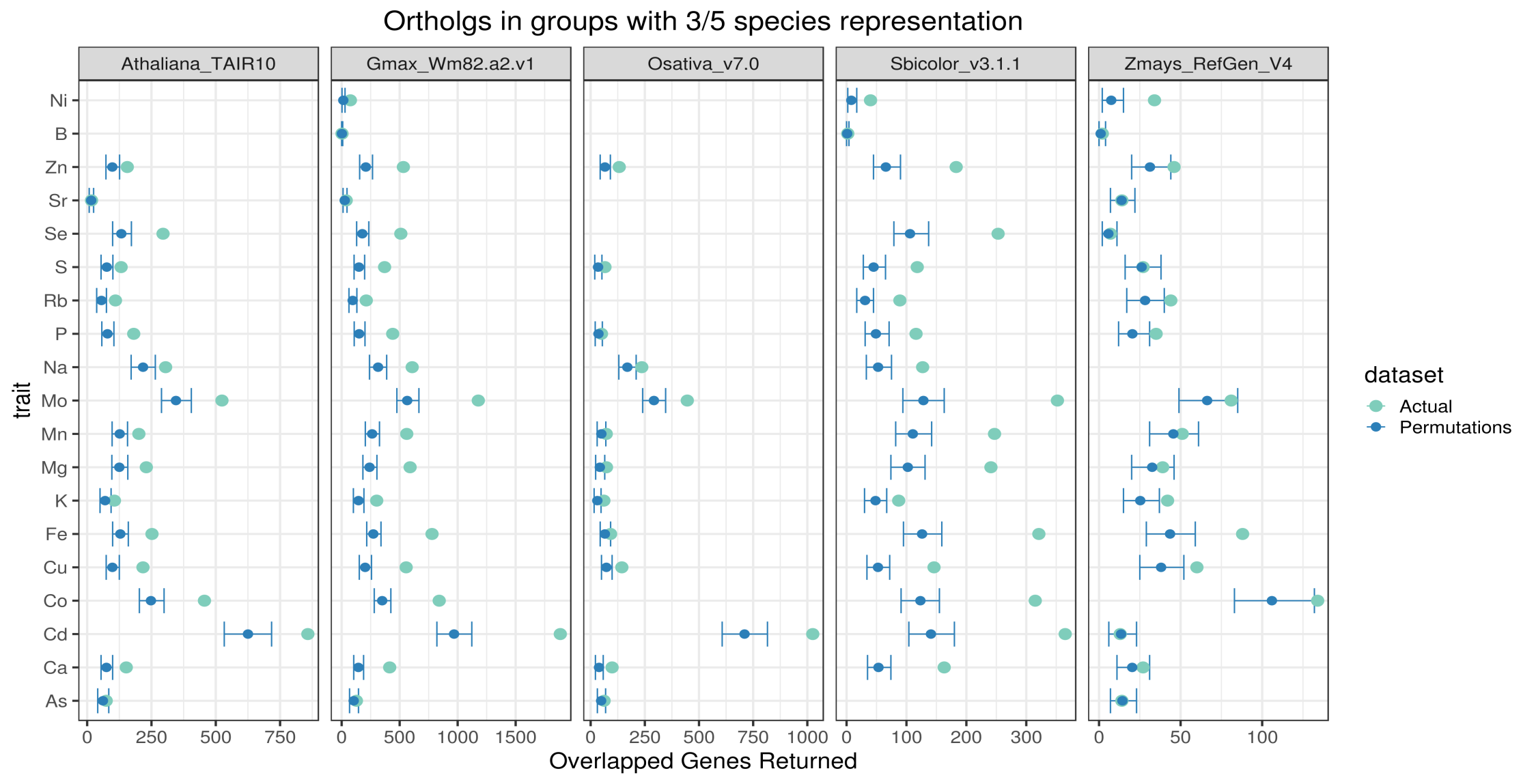

### S2_Fig.tiff

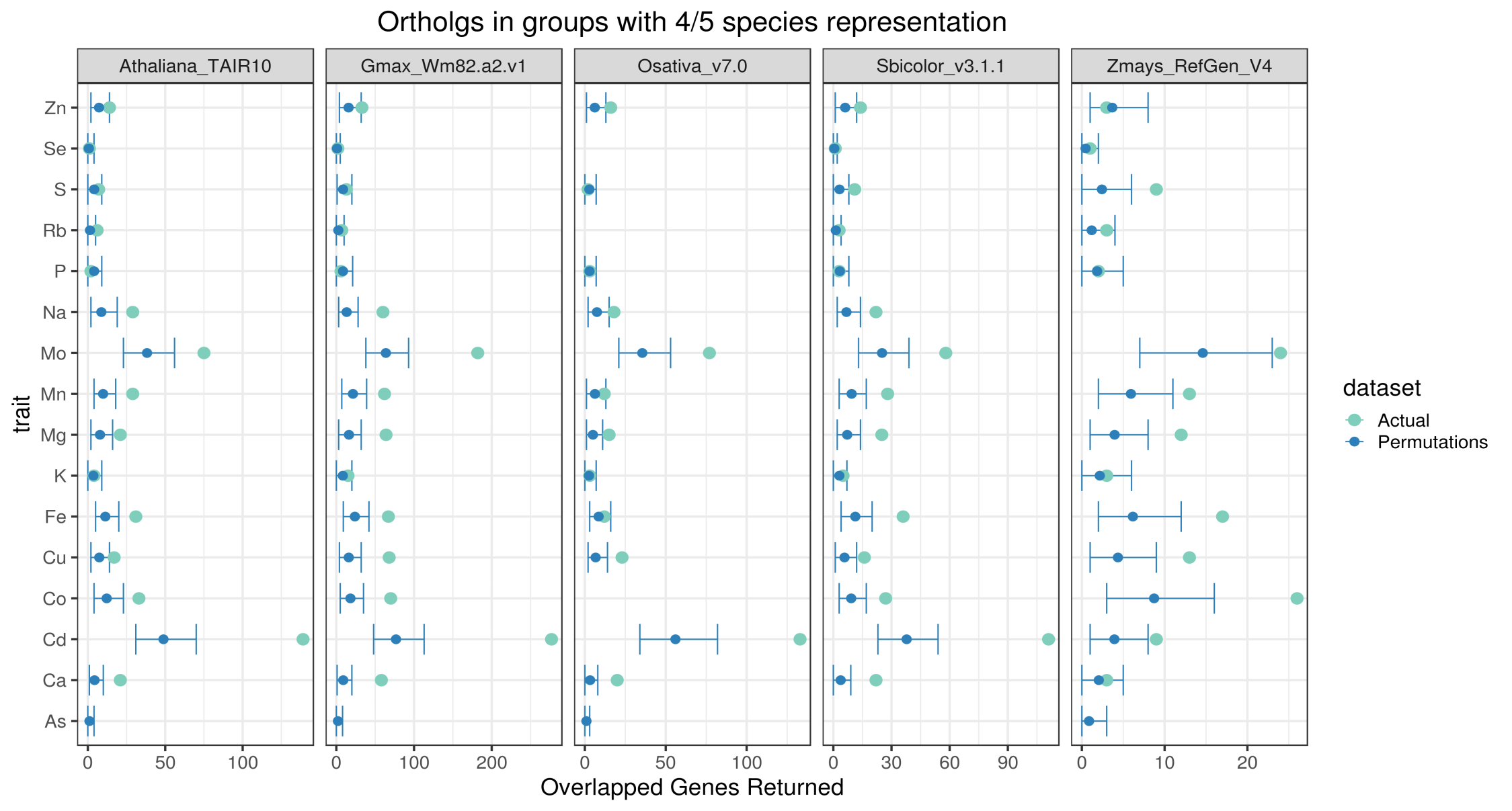

### S3_Fig.tiff

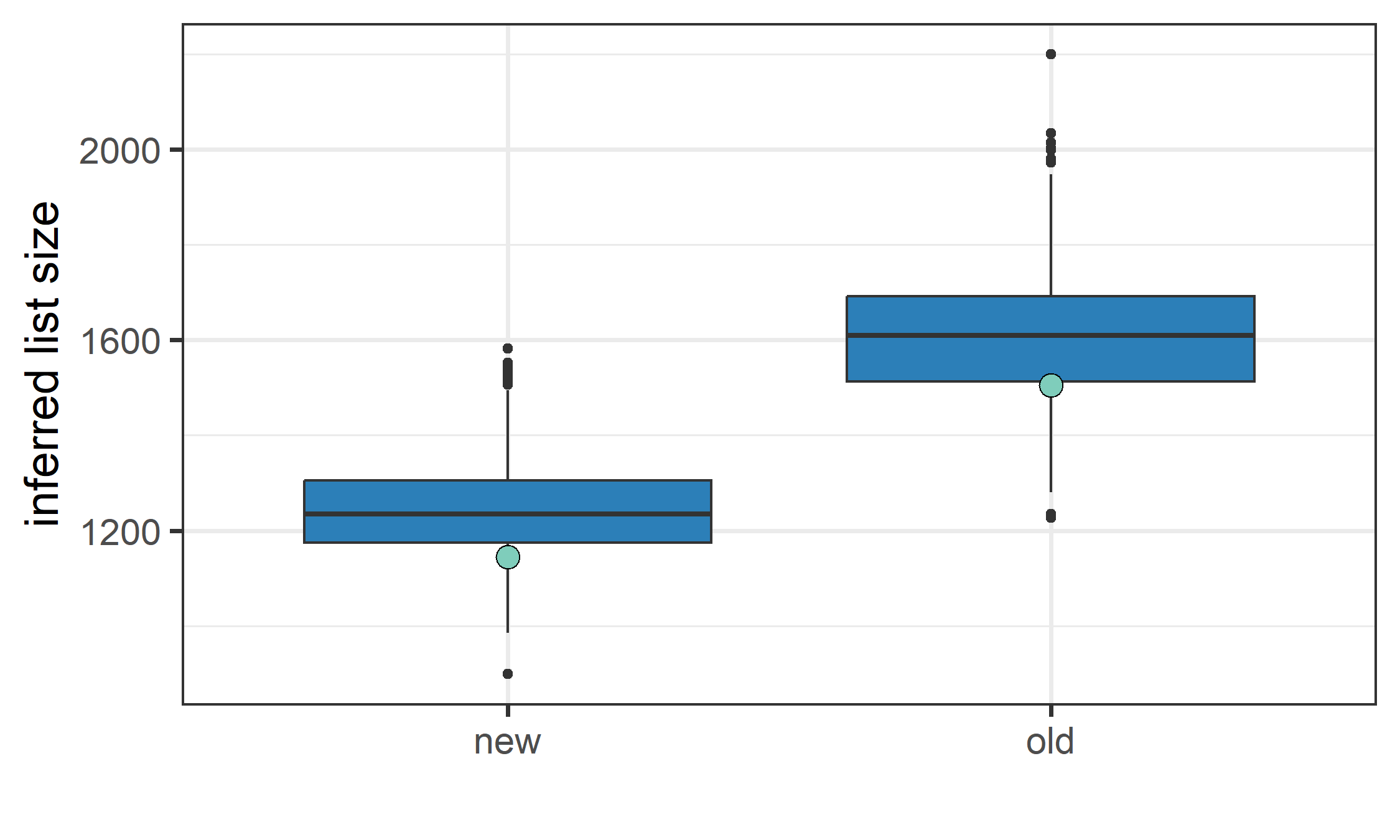

### S4_Fig.tiff

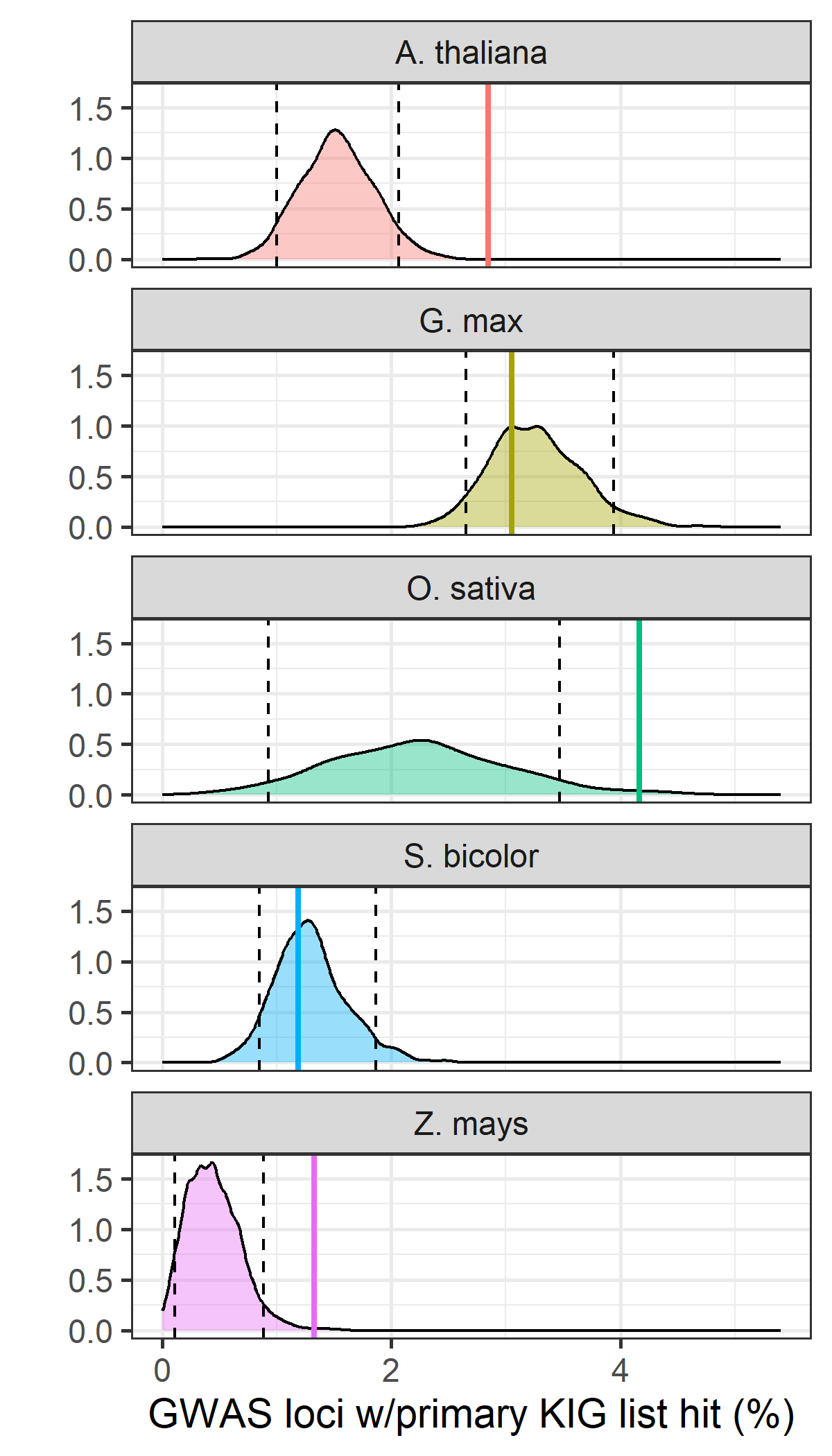

### S5_Fig.tiff

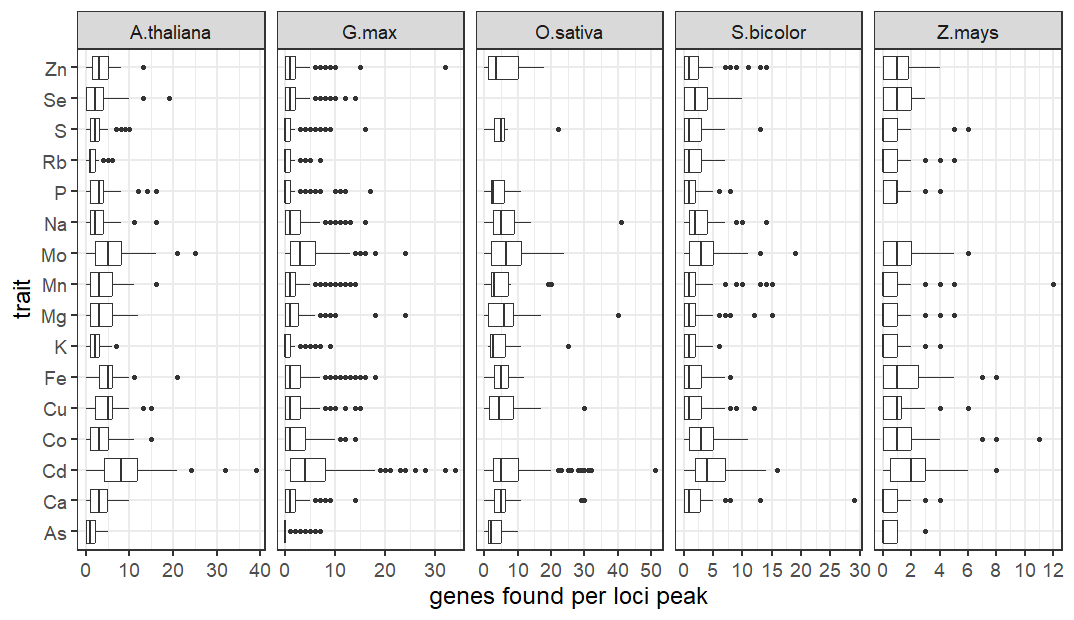
